# Seasonal Variability and Shared Molecular Signatures of Inactivated Influenza Vaccination in Young and Older Adults

**DOI:** 10.1101/719203

**Authors:** Stefan Avey, Subhasis Mohanty, Daniel G. Chawla, Hailong Meng, Thilinie Bandaranayake, Ikuyo Ueda, Heidi J. Zapata, Koonam Park, Tamara P. Blevins, Sui Tsang, Robert B. Belshe, Susan M. Kaech, Albert C. Shaw, Steven H. Kleinstein

## Abstract

The seasonal influenza vaccine is an important public health tool but is only effective in a subset of individuals. The identification of molecular signatures provides a mechanism to understand the drivers of vaccine-induced immunity. Most previously reported molecular signatures of influenza vaccination were derived from a single age group or season, ignoring the effects of immunosenescence or vaccine composition. Thus, it remains unclear how immune signatures of vaccine response change with age across multiple seasons. Here we profile the transcriptional landscape of young and older adults over five consecutive vaccination seasons to identify shared signatures of vaccine response as well as marked seasonal differences. Along with substantial variability in vaccine-induced signatures across seasons, we uncovered a common transcriptional signature 28 days post-vaccination in both young and older adults. However, gene expression patterns associated with vaccine-induced antibody responses were distinct in young and older adults; for example, increased expression of Killer Cell Lectin Like Receptor B1 (*KLRB1*; *CD161*) 28 days post-vaccination positively and negatively predicted vaccine-induced antibody responses in young and older adults, respectively. These findings contribute new insights for developing more effective influenza vaccines, particularly in older adults.

## Introduction

Influenza is a major public health burden, particularly in high-risk populations such as older adults. The seasonal inactivated influenza vaccination (IIV) is estimated to be 50-70% effective in randomized controlled trials of young adults (1–5), and efficacy is reduced to under 50% in adults over age 65 (6). Understanding the dynamics of vaccination-induced immune responses, and the factors associated with immunological protection should provide insights important for improving vaccine design.

Systems vaccinology approaches utilizing high-throughput immune profiling techniques have identified signatures of response to influenza vaccination (7–14). These include pre-vaccination transcriptional signatures of apoptosis-related gene modules (9), as well as B cell signaling and inflammatory modules (15). Post-vaccination transcriptional signatures have also been identified, including an early interferon response 1 day post-vaccination and a plasma cell response 3 and 7 days post-vaccination (13). Interferon stimulated genes were upregulated in both monocytes and neutrophils between 15 and 48 hours post-vaccination and correlated with influenza-specific antibody responses (7, 12). In addition, the expression of genes enriched for proliferation and immunoglobulin production 7 days post-vaccination accurately predicted antibody response in an independent cohort (10). Studies of the influence of aging revealed that an early interferon response 1-2 days post-vaccination as well as an oxidative phosphorylation and plasma cell response 7 days post-vaccination were correlated with antibody response in young adults but were diminished or dysregulated in older adults (13, 14).

Notably, previous studies of influenza vaccine response studying the effects of aging used data from a single vaccine season (9) or from two consecutive seasons in which vaccine composition was identical (13, 14); consequently, the generalizability of these signatures is unknown. To date, no comprehensive characterization of vaccine response in both young and older adults has been reported to multiple influenza vaccines which vary in composition. To address this gap, we profiled young and older adults over five consecutive vaccination seasons (2010-11, 2011-12, 2012-13, 2013-14, and 2014-15) hereafter referred to by the first year of each season. We developed a new automated metric to quantify antibody response while accounting for baseline titers and used this novel metric to identify predictive transcriptional signatures of vaccine response using post-vaccination as well as baseline gene expression profiles.

## Materials and Methods

### Clinical Study Design and Specimen Collection

A total of 317 subjects were recruited at Yale University over the five vaccination seasons between 2010 and 2014 and HAI titers pre- (D0) and post-vaccination (D28) were available from the 294 subjects reported in Table 1. Informed consent was obtained for all subjects under a protocol approved by the Human Subjects Research Protection Program of the Yale School of Medicine. Participants with an acute illness two weeks prior to recruitment were excluded from the study, as were individuals with primary or acquired immune-deficiency, use of immunomodulating medications including steroids or chemotherapy, a history of malignancy other than localized skin or prostate cancer, or a history of cirrhosis or renal failure requiring hemodialysis. Blood samples were collected into Vacutainer sodium heparin tubes and serum tubes (Becton Dickinson) at four different time points, immediately prior to administration of vaccine (D0) and on D2 (2011, 2012, 2013, 2014) or D4 (2010), D7, and D28 post-vaccination.

**Table 1.** Vaccine Compositions and Cohorts

|  | 2010-11 | 2011-12 | 2012-13 | 2013-14 | 2014-15 |
| --- | --- | --- | --- | --- | --- |
| **Vaccine Composition<sup>a</sup>** | A/California/7/2009 | A/California/7/2009 | A/California/7/2009 | A/California/7/2009 | A/California/7/2009 |
|  | A/Perth/16/2009 | A/Perth/16/2009 | A/Victoria/361/2011 | A/Texas/50/2012 | A/Texas/50/2012 |
|  | B/Brisbane/60/2008 | B/Brisbane/60/2008 | B/Wisconsin/1/2010 | B/Brisbane/60/2008 | B/Brisbane/60/2008 |
|  |  |  |  | B/Massachusetts/2/2012 | B/Massachusetts/2/2012 |
| **Subjects** | 42 | 69 | 92 | 56 | 35 |
| **Gender (% Male)** | 33 | 42 | 40 | 36 | 51 |
| **Age Group (% Older)** | 48 | 54 | 49 | 52 | 46 |
| **Transcriptomes<sup>b</sup>** | 19 | 39 | 30 | 26 | 20 |
| **Young (LR \| I \| HR)<sup>c</sup>** | 4 \| 1 \| 6 | 8 \| 2 \| 6 | 6 \| 0 \| 9 | 6 \| 2 \| 5 | 4 \| 2 \| 5 |
| **Older (LR \| I \| HR)<sup>c</sup>** | 5 \| 0 \| 3 | 11 \| 5 \| 7 | 7 \| 0 \| 8 | 7 \| 0 \| 6 | 2 \| 0 \| 7 |

<sup>a</sup> The three vaccine strains in 2009-10 were A/Brisbane/59/2007, A/Brisbane/10/2007, and B/Brisbane/60/2008. A monovalent A/California/7/2009 vaccine was administered to some subjects in March 2010.
<sup>b</sup> Subjects with transcriptional data are a subset of subjects with antibody titers.
<sup>c</sup> Subjects are listed by antibody response category: low responder (LR), indeterminate (I), high responder (HR).

In order to understand the transcriptional program underlying a successful vaccination response, we identified a subset of 134 subjects with extreme (strong or weak) antibody responses to perform transcriptional profiling by microarrays. In the first three seasons, the selection criteria were a four-fold increase to at least 2 strains (strong response) or no four-fold increase to any strain (weak response) as described previously (14). In the fourth and fifth seasons, the adjMFC metric was used in addition to the fold change criteria to account for baseline titers (11). The maxRBA response endpoint was developed after the study completed, however, less than 10% (12/134) of subjects chosen for transcriptional profiling had indeterminate responses by maxRBA (neither high or low responders using a 40% cutoff) (Table 1). These 12 subjects were excluded from the predictive modeling of antibody response.

### HAI and VNA Analyses, Cell Sorting, RNA processing and Gene Expression Analyses

Detailed methods are provided in *SI Appendix*.

## Results

### Antibody Titer Dynamics

We evaluated 294 healthy young (21 - 30 years old, n = 147) and older (≥ 65 years old, n = 147) adults over five consecutive influenza vaccination seasons from 2010-2014. All subjects received the standard dose trivalent (2010, 2011, 2012) or quadrivalent (2013, 2014) seasonal inactivated influenza vaccine (IIV). We measured influenza-specific hemagglutination inhibition (HAI) titers pre-vaccination (D0) and 28 days post-vaccination (D28). Over the course of our study, the vaccine composition changed relative to the previous season in three of five seasons (Table 1).

In all seasons, pre-vaccination titers were negatively correlated with the increase in titers post-vaccination (*SI Appendix*, Fig. S1). Previous work defined an adjusted maximum fold change (adjMFC) endpoint that removes the nonlinear correlation between fold change and baseline titers (11). However, adjMFC separates subjects into manually defined bins, making it difficult to perform high-throughput analysis. Furthermore, adjMFC does not allow for information sharing between bins as each bin is adjusted independently. To address these limitations, we developed maximum Residual after Baseline Adjustment (maxRBA), which corrects for the dependence on baseline titers for each strain by modeling titer fold changes as an exponential function of pre-vaccination titers and selecting the maximum residual across strains (Fig. 1A). All vaccine strains were approximately equally responsible for the maximum residual in any given season. “High responders” (HR) and “low responders” (LR) were defined as the top and bottom 40th percentiles of the residuals, respectively. maxRBA can be interpreted as the maximum change from expected fold change given the initial titer; it is fully automated, is strain agnostic, and is correlated with plasmablast frequencies seven days post-vaccination (*SI Appendix*, Fig. S2A-B). Thus, maxRBA allows a completely automated assessment of the relative strength of each subject’s antibody response independent of pre-existing antibody titers.

**Figure 1.**
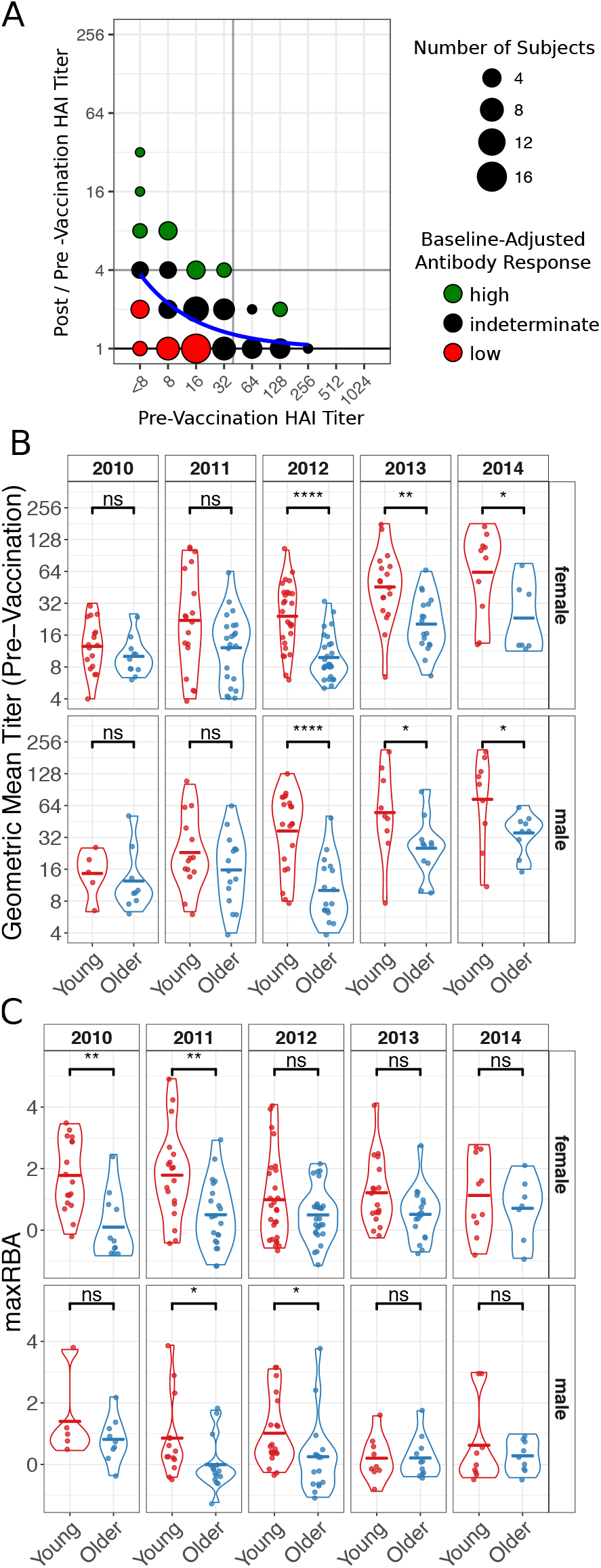
Influenza-Specific Antibody Titers. (A) An illustration of the maximum Residual after Baseline Adjustment (maxRBA) method for hemagglutination inhibition (HAI) titers to the B/Wisconsin/1/2010 strain in the 2012 season. An exponential curve (blue) is fit to the data and the residual is used to stratify subjects into high and low responders. Subjects with largest positive residuals are high responders (green) and subjects with smallest negative residuals are low responders (red). maxRBA is calculated using the maximum residual across all vaccine strains. (B and C) Violin plots of pre-vaccination HAI titers (B) and HAI responses measured by maxRBA (C) are separated by season and gender to compare age groups. Crossbars indicate the mean. Not Significant (ns) p > 0.05, * p < 0.05, ** p < 0.01, *** p < 0.001, **** p < 0.0001 independent two-sided Wilcoxon rank sum test.

Older adults had significantly lower pre-vaccination titers than young adults for three of five seasons (Fig. 1B). The maximum fold change to any vaccine strain showed an increasing trend in young adults compared to older adults (*SI Appendix*, Fig. S3C). Because of the inverse relationship between baseline titers and fold change (*SI Appendix*, Fig. S1), we adjusted for baseline titers using maxRBA and found that the difference in vaccine response between young and older adults was statistically significant in more seasons (Fig. 1C). Males and females had similar pre-vaccine geometric mean titers (preGMTs) (*SI Appendix*, Fig. S3A). However, the antibody response calculated by maxRBA showed a trend toward stronger antibody responses in females compared to males with similar baseline titers in both age groups (Fisher’s combined p = 0.02 (Young), p = 0.12 (Older); *SI Appendix*, Fig. S3B). We did not detect any significant difference in baseline titers or titer responses across seasons when stratifying subjects by body mass index, smoking history, aspirin use, or diabetes medication use (p > 0.05 two-sided Wilcoxon rank sum test (discrete) or simple linear regression (continuous)).

We also examined the dynamics of viral titers over the course of the five seasons (*SI Appendix*, Fig. S3D). The A/California 7/2009 H1N1 strain was introduced into the seasonal vaccine in 2010 and remained through the 2014 season; however, pre-vaccine titers to this strain were consistently lower in older vs. young adults for 2011-2014. While we did not follow the same subjects across multiple seasons, 50-80% of young and 80-98% of older adults self-reported receiving influenza vaccine in the previous year. Taken together, these results support existing evidence that the capability for antibody persistence is reduced with age (16).

### Substantial Seasonal Variability in Vaccine-Induced Signatures

To identify correlates and predictors of vaccine response, we selected a subset of individuals (20 - 40 subjects per season) from young and older adult cohorts who had strong or weak antibody responses according to HAI titers and performed longitudinal transcriptional profiling pre-vaccination (baseline) and 4 (2010 cohort) or 2 days (all other cohorts), 7 days, and 28 days post-vaccination (Table 1; *Methods*). We first performed differential expression analysis independently in each season without differentiating subjects by antibody response. We compared each post-vaccination time point to baseline and found a vaccine-induced signature that comprised a total of 2,462 significantly differentially expressed genes (DEGs) over all five seasons (FDR < 0.05, Fold Change > 1.25; *SI File 1*).

Most of the DEGs were from the first two seasons whereas vaccination in the latter three seasons induced relatively weak changes (Fig. 2A; *SI Appendix*, Fig. S4E). In fact, a substantial fraction of DEGs were unique to a single season and not differentially expressed at any time point in another season (Young: 38%, Older: 75%). In young adults, there were 1,330 DEGs shared across two or more seasons while in older adults there were 265 shared DEGs. In both young and older adults, a substantial fraction of these shared genes was differentially expressed 28 days post-vaccination (*SI Appendix*, Fig. S4F). To assess whether vaccine-induced changes were consistent between seasons, we divided the 2,462 DEGs into 7 clusters by hierarchical clustering (Fig. 2A; *SI File 2*) and tested for their activity in every season using QuSAGE (17) (*SI Appendix*, Fig. S5). In young adults, three of the clusters (B, F, G) had significant, but opposite, activity during the 2010 and 2011 seasons, while these clusters were relatively consistent across seasons in older adults. Genes in cluster A were induced strongly in the 2011 season in both age groups and notably enriched for multiple pathways related to mitochondria, including *mitochondrial inner membrane, oxidative phosphorylation, respiratory electron transport, citric acid (TCA) cycle and respiratory electron transport*, and *mitochondrial respiratory chain complex assembly* (FDR < 0.05; *SI File 2*). These findings reflect our previous identification of a mitochondrial biogenesis signature associated with influenza vaccine antibody response (14). Cluster D was only significantly induced in the 2013 season at 7 and 28 days post-vaccination and was not significantly enriched for any gene sets tested (FDR > 0.05; *SI File 2*). The cluster with the most consistent expression pattern across the five seasons was cluster C, which was enriched for pathways related to Toll-like receptor signaling, B and T cell signaling, NF-κB signaling, MAPK signaling, cell senescence or proliferation, and apoptosis (*SI File 2*). Interestingly, cluster C contains three genes (*DUSP1, DUSP2, CCL3L3*) which were significantly downregulated 28 days post-vaccination in four of five seasons. *CCL3L3* is a ligand for *CCR1, CCR3* and *CCR5*, known to be chemotactic for monocytes and lymphocytes (18). *DUSP1* and *DUSP2* are dual specificity phosphatases; *DUSP2* dephosphorylates *STAT3*, leading to inhibition of survival and proliferation signals (19–21), and an age-associated decrease in *DUSP1* function contributed to inappropriate IL-10 production in monocytes before and after influenza vaccination (22). To determine whether downregulation of these three genes was a result of changes in cell subset composition or observed in subpopulations of cells, we performed transcriptional profiling on sorted B and T cells in a subset of individuals from three seasons. *DUSP1* and *DUSP2*, but not *CCL3L3*, were significantly downregulated 28 days post-vaccination over multiple seasons in CD4 and CD8 T cells of young adults (One sided t-test p < 0.01; Fig. 2C, *SI Appendix*, Fig. S4C-D). Furthermore, while *DUSP2* was only significantly decreased in PBMCs of older individuals in the 2011 season, expression of *DUSP2* was significantly decreased 28 days post-vaccination in sorted CD4 and CD8 T cells from older individuals in multiple seasons (Fig. 2C). Thus, the downregulation of *DUSP2* 28 days post-vaccination is observed in the T cell compartment of both young and older adults.

**Figure 2.**
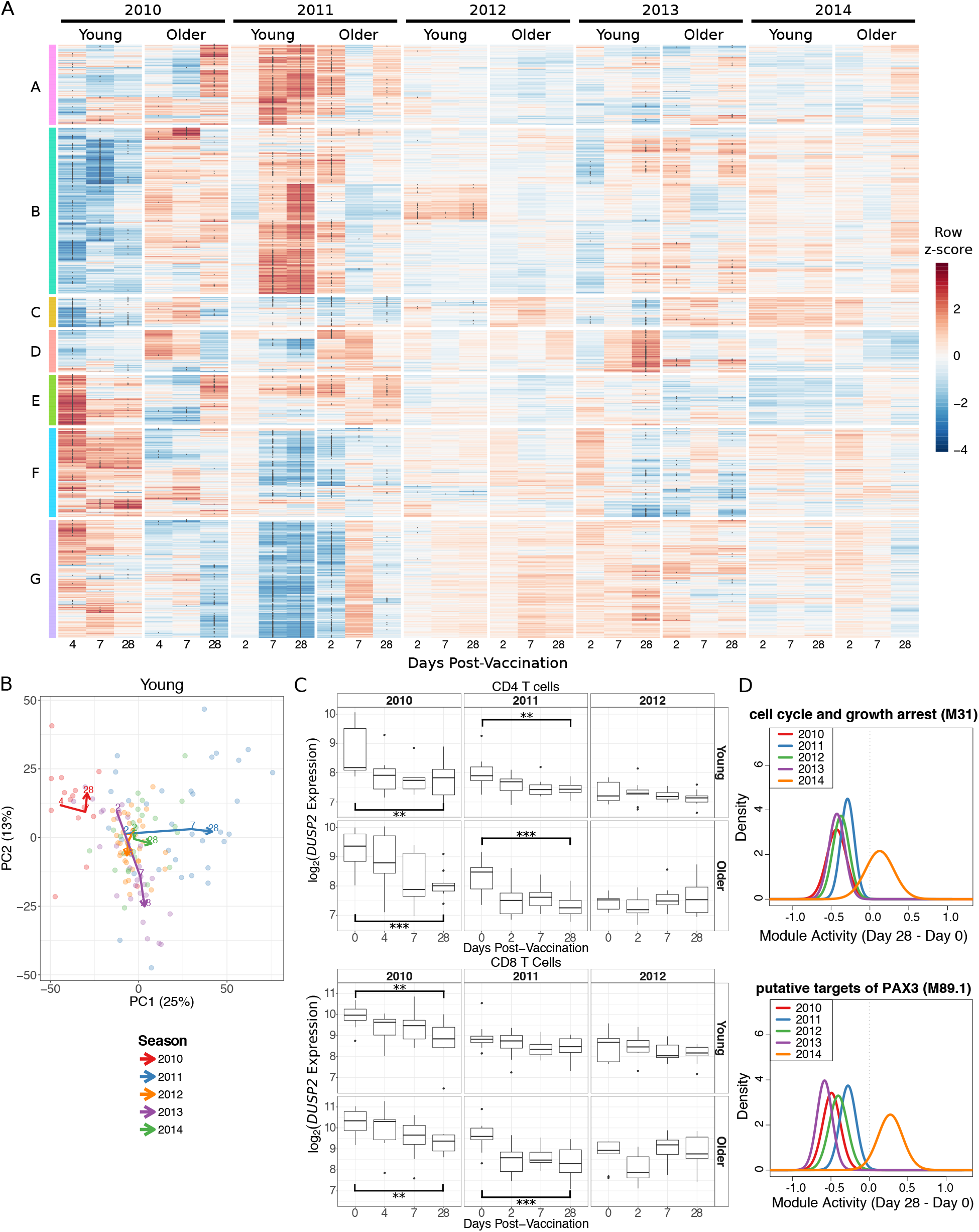
Substantial Seasonal Variability in Signatures Induced by Influenza Vaccination. (A) A row-normalized heatmap of the 2,462 significantly differentially expressed genes (DEGs). Clusters A-G were defined by hierarchical clustering. Asterisks within the heatmap indicate genes significantly differentially expressed compared to day 0. (B) The first two principal components from a principal component analysis of all DEGs. Each point is a sample and lines connect the median of the points at each day post-vaccination within each season. (E) *DUSP2* expression in sorted CD4 and CD8 T cells. ** p < 0.01, *** p < 0.001 one-sided t-test comparing day 28 and day 0 only. (F) Probability density functions calculated by QuSAGE for two representative gene modules significantly downregulated 28 days post-vaccination in four seasons. M31 contains *DUSP1* while M89.1 contains both *DUSP1* and *DUSP2*.

To further assess shared patterns in vaccine-induced changes across five seasons, we performed a principal component analysis (PCA) on gene expression fold changes post-vaccination for all DEGs. The first two components together explained 38% of the variation in young adults’ and 46% of the variation in older adults’ transcriptional changes post-vaccination (Fig. 2B, *SI Appendix*, Fig. S4B). Notably, in young adults, the 2011 and 2014 seasons (both with vaccine composition identical to the previous year) had similar trajectories, increasing along the first principal component (PC1) by D28 post-vaccine. Examining the genes contributing to PC1 reveals that four of the top 10 genes (*SLMAP, MATR3, MBNL3, RANBP3*) increase in expression post-vaccination more in the 2011 and 2014 seasons than in any other season. The shared trajectories along PC1 are not significantly enriched for any blood transcription modules (BTMs) (23), KEGG pathways (24), or cell subset signatures (25) (FDR > 0.05; *SI File 3*). The trajectory of the 2010 season was quite distinct from the other seasons in young adults. This season is consistently elevated on PC2, which is significantly enriched for monocytes, TLRs and inflammatory signaling (FDR < 0.05; *SI File 3*). The 2012 and 2013 seasons also appear to have similar trajectories, both decreasing in PC2 over time. The vaccines used in these two seasons each introduced multiple new strains while retaining the A/California/7/2009 strain. Five of the top 10 genes (*ZNF493, ZNF652, OCIAD1, C21orf58, IL11RA*) contributing to PC2 increased in expression 28 days post-vaccination in the 2012 and 2013 seasons while decreasing in expression in the other seasons. This differential expression analysis shows that there are large variations in vaccine-induced transcriptional signatures between seasons which, in young adults, might be explained in part by vaccine composition.

Given the substantial seasonal variation in the number of DEGs, we next performed an analysis of differential expression of gene modules using QuSAGE to quantify the gene module activity of 346 previously defined BTMs (23). There were 262 differentially expressed modules (DEMs) (FDR < 0.05; *SI File 4, SI Appendix*, Fig. S4A). Similar to the gene-level analysis, no significant changes were identified in the 2014 season, but six modules (*cell cycle and growth arrest (M31), chemokines and inflammatory molecules in myeloid cells (M86.0), enriched for TF motif TTCNRGNNNNTTC, leukocyte differentiation (M160), putative targets of PAX3 (M89.1), and signaling in T cells (I) (M35.0)*) were significantly downregulated in young adults at D28 in four of five seasons (Fig. 2D). These changes were largely driven by decreases in *DUSP1/2, EGR1/2, JUN/JUNB, FOS/FOSB, TNF, CD83*, and *IL1B*. Thus, while there was substantial variability in the signatures induced by vaccination across multiple seasons, there is a shared signature consisting of three genes and six modules which was downregulated at D28 in four of five seasons.

### Shared Vaccine-Induced Signatures Across Five Seasons

The differential expression approach is limited by fixed fold change and significance cutoffs that may vary between seasons. To increase our power to identify shared signatures across seasons and in older adults, we performed a meta-analysis at the individual gene and gene module level. We identified 338 genes with significantly altered expression post-vaccination (FDR < 0.05; *SI File 5*). In young adults, we identified significant genes at D2, D7 and D28 with little overlap among genes on each day. Genes induced on D2 were moderately enriched for innate immune genes from InnateDB (http://innatedb.com/) including *MYH9, TYK2, GLRX*, and *IP6K1* (p = 0.12, hypergeometric test). Some of the genes consistently induced at D7 included *IGLL1, CD38, ITM2C, TNFRSF17, MZB1*, and *TXNDC5*. We previously identified *TNFRSF17*, B cell maturation antigen, as induced seven days following influenza vaccination (26), and it was also identified as a predictive marker gene of antibody response to multiple vaccines including influenza, meningococcal conjugate (MCV4), and yellow fever (YF17D) vaccines (11, 23, 27–29). Consistent with the individual season analysis, the majority of genes identified by the meta-analysis were altered at D28; these D28 DEGs included *DUSP1, DUSP2*, and *CCL3L3*, identified in the single-season analysis, and many other downregulated genes including *IL1B, CCL3*, and *JAK1*. Thus, there are consistent changes identified across all seasons in young adults at every time point measured.

In older adults, we identified 125 genes with significantly altered expression at D28, but no genes with significantly altered expression at D2 or D7 (*SI File 5*). The most significantly increased gene at D28 is *XRN1*, the primary 5’ to 3’ cytoplasmic exonuclease involved in mRNA degradation (30). *XRN1* plays a critical role in the control of RNA stability in general, but in addition appears to regulate the response to viral infection at several levels—for example, by targeting viral RNAs for degradation (31), or regulating levels of potential activating ligands such as double-stranded RNA (32). Notably, *XRN1* has also been reported to facilitate replication of influenza and other viruses by inhibiting host gene expression (33, 34) - suggesting that dysregulated expression of *XRN1* in older adults could influence host response to vaccination. We identified 3 genes shared between both age groups: *ARRDC3* and *USP30* were downregulated while *TNPO1* was upregulated, all at D28. *ARRDC3* encodes a member of the arrestin protein family which regulates G protein-mediated signaling and is implicated in regulating metabolism (35). *USP30* is a ubiquitin-specific protease that acts as a mitochondrial deubiquitinating enzyme (36). *TNPO1* encodes Transportin-1 that serves to import proteins into the nucleus (37). The effect sizes of all genes at D28 were positively correlated between young and older adults with weak positive associations at D2 and D7 (Fig. 3). These results provide additional evidence that transcriptional changes are broadly similar in young and older adults at D28 post-vaccine.

**Figure 3.**
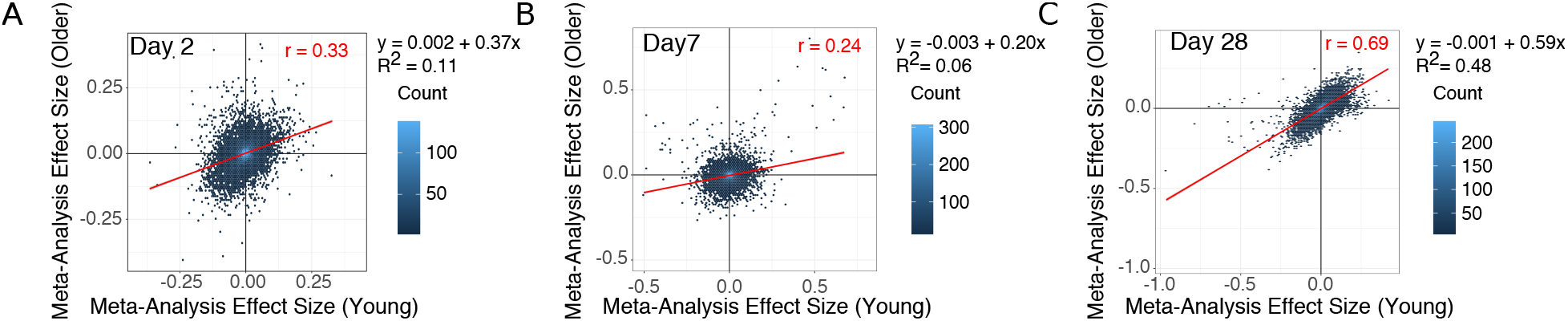
Vaccine-Induced Changes are Correlated Between Young and Older Adults at Day 28. Scatter plots show the meta-analysis effect sizes of changes post-vaccination for every gene in young vs older adults on days 2 (A), 7 (B) and 28 (C) post-vaccination.

We carried out a gene set level meta-analysis using QuSAGE to combine probability density estimates of gene module activity for each season (38). We identified 186 BTMs significantly altered post-vaccination across five seasons (FDR < 0.05; *SI File 4*). The module with the largest increase in activity was *plasma cells, immunoglobulins (M156.1)* which peaked on D7 with a combined fold change of 1.17 in young adults and 1.08 in older adults at D7. Most BTMs showing significant changes were identified in young adults and, unlike the individual gene level, there was a large overlap between sets at each time point, suggesting the same module changes were sustained over the 28 days following vaccination (*SI Appendix*, Fig. S4A). Indeed, a heatmap of module activity shows that in young adults, transcriptional changes continued to intensify at D28 for many modules rather than returning to the baseline state (*SI Appendix*, Fig. S6). Older adults showed a qualitatively similar pattern to young adults on D2 and D28, but not D7. The majority (40/59) of the modules significantly altered in older adults on D28 were also significantly altered in young adults at D28 (*SI Appendix*, Fig. S4A). The modules downregulated on D28 in both young and older adults were annotated with antigen processing and presentation (M95.0, M95.1, M28, M71, M200, M5.0) and T cell activation (M36, M44, M52). The modules upregulated on D28 included *golgi membrane (II) (M237), enriched in DNA interacting proteins (M182)*, and *chaperonin mediated protein folding (I, II) (M204.0, M204.1*). Taken together, the high correlation between individual gene changes and overlap of many BTMs suggest a convergence toward a common transcriptional program in young and older adults at D28.

### Age-Associated Genes are Induced 7 Days Post-Vaccination

A meta-analysis across all five seasons revealed markedly different baseline transcriptional profiles in young vs. older adults, with 1,072 genes significantly altered (FDR < 0.05, *SI File 6*). Of these age-associated genes, 204 genes were also significantly induced by the vaccine in young adults and 125 genes in older adults. We tested whether age-associated genes were enriched for vaccine-induced genes at each time point and found that the overlap was significantly more than expected by chance for the 6 age-associated genes induced on D7 in young adults (p = 0.017, hypergeometric test). Of these 6 overlapping genes, 5 genes (*ITM2C, MZB1, IGLL1, TNFRSF17*, and *TXNDC5*) exhibited decreased basal expression in older adults while 1 (*SELENOS*) exhibited increased basal expression compared to young adults. While these genes were induced in young adults, they were not significantly induced in older adults on D7. Notably, *MZB1* and *TNFRSF17* are B cell associated genes, suggesting that older adults have decreased B cell activity pre-vaccination and fail to induce the same B cell response as young adults at D7. *SELENOS* encodes selenoprotein S, which is involved in degrading misfolded endoplasmic reticulum (ER) proteins and influences inflammation via the ER stress response (39, 40). Our results show that age-associated genes are significantly over-represented in the set of genes altered in young adults 7 days post-vaccination.

We next performed a meta-analysis of BTMs between age groups at baseline and identified 120 modules significantly altered with age (FDR < 0.05, *SI File 7*). Most of the modules that were decreased with age were associated with adaptive immunity, whereas those that had increased expression with age were mostly innate and inflammatory modules (reflecting age-associated inflammatory dysregulation; *SI Appendix*, Fig. S7B). Of these 120 modules, 52 were also significantly altered post-vaccination; however, the overlap at each time point was not significantly more than expected by chance (hypergeometric p > 0.05, *SI Appendix*, Fig. S7A). Thus, age-related genes are enriched among the genes induced at D7 in young adults while no gene modules were significantly over-represented.

### Post-Vaccination Predictors of Antibody Response

We next asked whether any transcriptional changes post-vaccination could discriminate high antibody responders (HR) from low antibody responders (LR). Regularized logistic regression models with an L1 (Lasso) or L1 and L2 (Elastic Net) penalties were fit to identify genes predictive of antibody response. In addition, to identify biologically interpretable predictors we used the Logistic Multiple Network-constrained Regression (LogMiNeR) framework (26) that facilitates the generation of predictive models with improved biological interpretability over standard methods. We combined the fold changes in gene expression data post-vaccination from five seasons and trained LogMiNeR to predict HR vs. LR in young and older cohorts separately (*SI Appendix*). At each time point, models were trained on all five seasons of data (except for D2, which was not available in the 2010 season; see *Methods*). Publicly-available data sets from independent groups were used to validate the models. For the models built from expression changes at D2 or D28, no studies at identical time-points were available, so we attempted to validate these models on studies with similar time points (day 1 or 3 in (11) and day 14 in (13)). While we could build predictive models on our data (median AUC ≥ 0.75) they did not validate on other data sets at the (different) time points available (median AUC ≤ 0.55).

For D7 post-vaccine, direct validation data were available in independent datasets. In young adults, D7 models were predictive for HR in the discovery and validation (11) datasets (Fig. 4A). Another MAP kinase phosphatase acting on ERK1/2, *DUSP5*, was one of 37 genes selected by the Lasso model whose expression was increased in HR (Fig. 4C). *DUSP5* is expressed in multiple immune cell types such as B cells (including plasma cells), T cells, dendritic cells, macrophages and eosinophils (41). In murine T cells, DUSP5 appears to promote the development of short-lived effector CD8+ T cells and inhibit memory precursor effector cell generation in an LCMV infection model (42); while optimizing memory precursor cell generation would be the goal of vaccination, the upregulation of *DUSP5* in HR could reflect regulation of the balance between short-lived vs. memory precursor effector CD8+ T cells. A sensitivity analysis of the maxRBA cutoff shows that the average expression of predictive genes is consistent across a range of definitions for HR and LR (20^th^ – 40^th^ percentile; *SI Appendix*, Fig. S2C-D). Using LogMiNeR, the models were consistently enriched for the *B Cell* signature as well as the KEGG *chemokine signaling pathway* (*SI File 8*).

**Figure 4.**
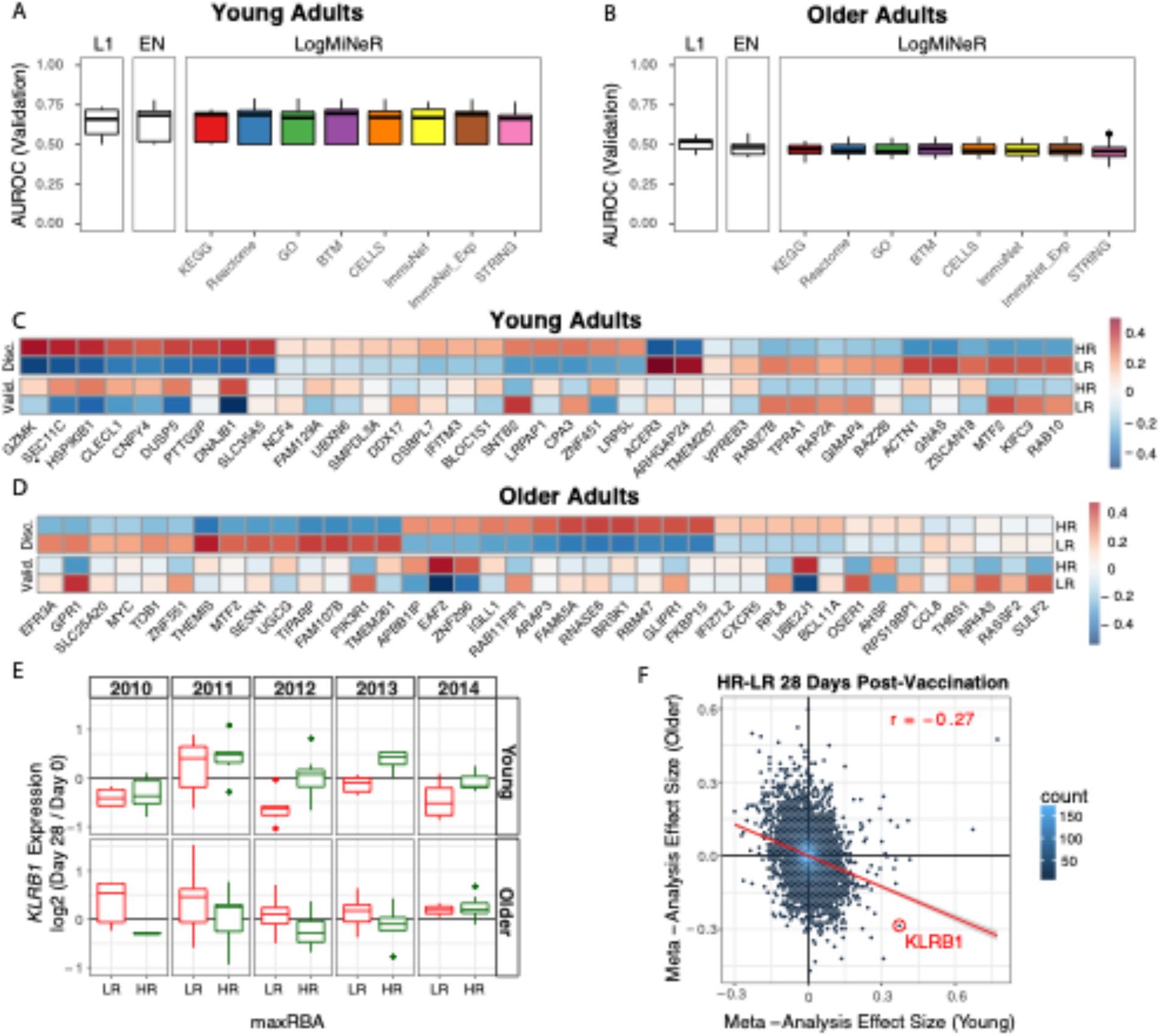
Post-Vaccination Transcriptional Predictors of Antibody Response. (A and B) Boxplots of the area under the receiver operating characteristic curve (AUROC) in the validation data for Lasso (L1), Elastic Net (EN), and Logistic Multiple Network-constrained Regression (LogMiNeR) models built from day 7 transcriptional changes in young (A) and older (B) adults. 50 iterations of cross-validation were performed. x-labels indicate the prior knowledge network for LogMiNeR (see *SI Appendix*). (C and D) Heatmaps of Discovery (Disc.) and Validation (Valid.) data showing the z-score of the fold change for individual genes selected by the L1 models in any iteration for young (C) and older (D) adults. (E) Boxplots of *KLRB1* expression changes in PBMCs 28 days post-vaccination in low responders (LR) and high responders (HR). (F) A scatter plot of the gene effect sizes comparing HR to LR 28 days post-vaccination in young vs older adults. *KLRB1* is indicated as a gene that has a positive effect size in one age group and negative effect size in the other.

In older adults, models predicting antibody responses built from D7 gene expression were highly predictive in the discovery dataset but did not validate on an independent dataset (13) (Fig. 4B, D). Expression of the Solute Carrier Family 25 gene *SLC25A20* of mitochondrial transporters contribute to predicting HR vs. LR in older adults. SLC25A20 is the carrier for carnitine and acylcarinitine (43), and so would be expected to be crucial for the transport of fatty acids into mitochondria. The models of response in older adults were significantly enriched for several BTMs of monocyte signatures as well as *TLR and Inflammatory Signaling* (*M16*), which positively predicted vaccine response; together with previous studies linking age-associated impairments in TLR function to influenza vaccine antibody response (44, 45), these findings provide additional support for the crucial role of innate immune function in vaccination (*SI File 8*).

Notably, none of the models built in young adults at any time point are predictive in older adults (AUC ≤ 0.5). In fact, models built on transcriptional changes at D28 in young adults had a median AUC near 0.8 in young adults, but no more than 0.3 in older adults, suggesting that the same genes predictive of HR in young adults predicted LR in older adults (*SI Appendix*; Fig. S8E). The Lasso models making these predictions often chose a single gene, Killer Cell Lectin Like Receptor B1 (*KLRB1*, also known as *CD161*), which was driving this inverse pattern (Fig. 4E). *KLRB1* is an inhibitory receptor on NK cells (46, 47) and is also a biomarker of Th17 cells (48–50). Notably, changes in *KLRB1* expression in sorted CD4 and CD8 T cells at D28 closely mirrored the changes in PBMCs for young, but not older adults (*SI Appendix*, Fig. S8A-B). We confirmed this inverse correlation between age groups on a genome-wide scale by performing a meta-analysis comparing HR vs. LR (*SI File 9*). We observed a weak negative correlation in effect sizes between young and older adults at D28 (r = −0.27; Fig. 4F). We confirmed this negative correlation in effect sizes between young and older adults using a virus neutralization assay (VNA) in a test sample of blood from seasons 2011 and 2012 (r = −0.32; *SI Appendix*, Fig. S8D). Thus, expression changes of many genes at D28 have opposing signs between age groups for the effect size comparing HR vs. LR, and a single gene, *KLRB1*, predicts response with AUC > 0.7 in opposing directions in young vs. older adults.

### Baseline Predictors of Antibody Response

We next sought to identify baseline transcriptional predictors of antibody response. In young adults, LogMiNeR models were predictive above random on discovery and validation (11) (Fig. 5A) datasets. Lasso models included the gene *VASH1*, known as an angiogenesis inhibitor and mediator of stress resistance in endothelial cells, which was expressed at lower levels in HRs (Fig. 5C); notably, the KEGG gene set *leukocyte transendothelial migration* was significantly enriched in over 50% of the models when LogMiNeR was used with ImmuNet as prior knowledge (51). Another predictive gene, *EIF4E*, a translation initiation factor important in type I interferon production, was decreased in HRs. A sensitivity analysis of the maxRBA cutoff shows that the average expression of predictive genes is consistent across a range of definitions for HR and LR (20^th^ – 40^th^ percentile; *SI Appendix*, Fig. S2E-F). Finally, the BTMs *cell adhesion (M51)* and *B cell surface signature (S2)* were consistently enriched in the models (*SI File 8*). In older adults, LogMiNeR models were also predictive on the discovery and one validation dataset (9) (Fig. 5B) but not another (13) (*SI Appendix*, Fig. S8C). Two of the individual genes that predict response, *ALDH1A1* and *ALDH3B1*, are aldehyde dehydrogenases which metabolize vitamin A to retinoic acid (Fig. 5D). Recently, aldehyde dehydrogenases were implicated in antiviral innate immunity as mediators of the interferon response through their role in the biogenesis of retinoic acid (52). Multiple monocyte gene sets are enriched in the predictive genes, including the BTM *enriched in monocytes (II) (M11.0)*, which negatively predicts vaccine response (*SI File 8*). Thus, these baseline predictive models built from five seasons of transcriptional profiling data provide further evidence for functional distinctions present in subjects prior to vaccination that influence the immunologic response to influenza vaccine in young and older adults.

**Figure 5.**
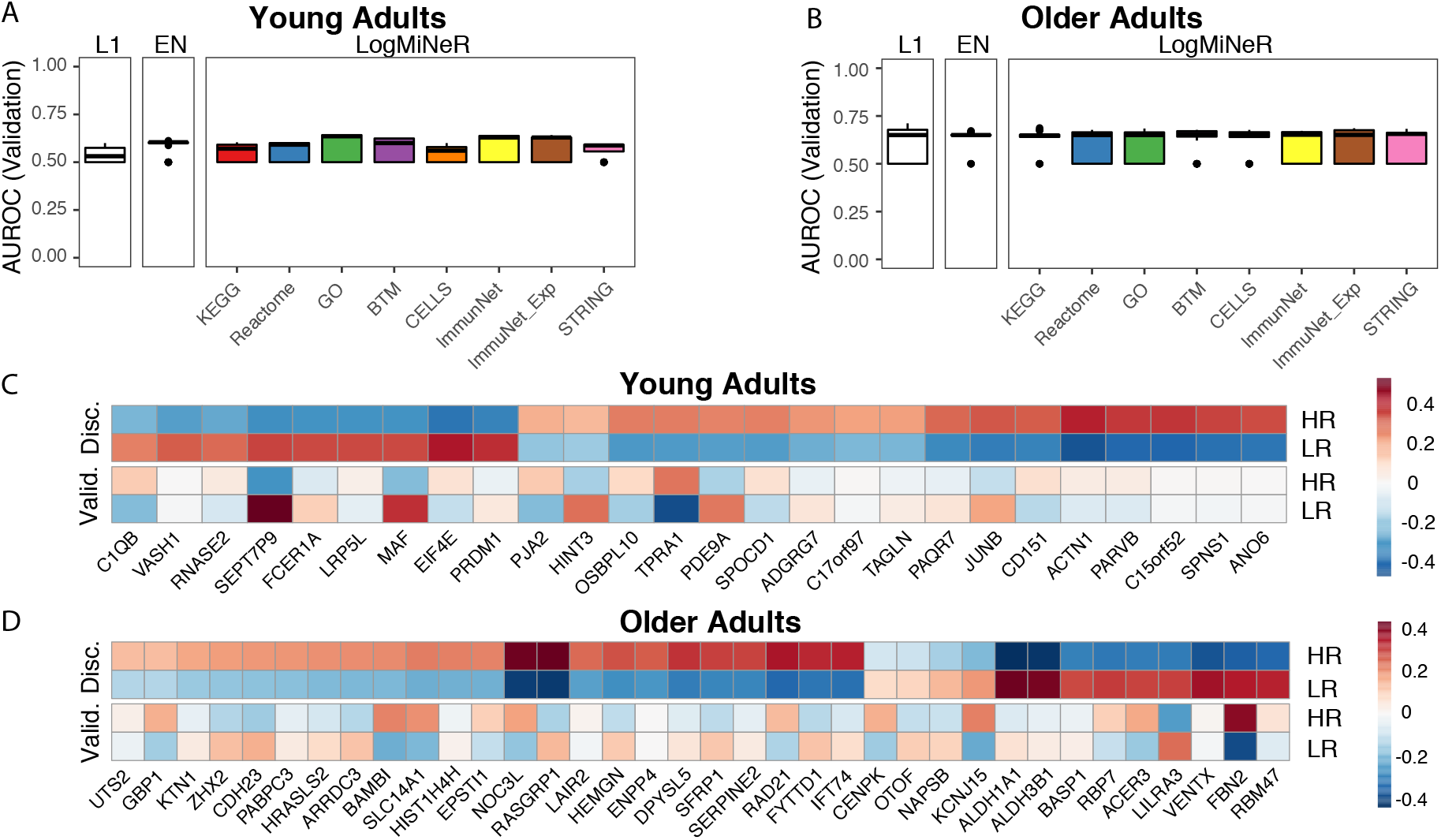
Baseline Transcriptional Predictors of Antibody Response. (A and B) Boxplots of the area under the receiver operating characteristic curve (AUROC) in the validation data for Lasso (L1), Elastic Net (EN),and Logistic Multiple Network-constrained Regression (LogMiNeR) models built from baseline (pre-vaccination) transcriptional profiles in young (A) and older (B) adults (9). 50 iterations of cross-validation were performed. x-labels indicate the prior knowledge network for LogMiNeR (see *SI Appendix*). (C and D) Heatmaps of Discovery (Disc.) and Validation (Valid.) data showing the z-score of the fold change for individual genes selected by the L1 models in any iteration for young (C) and older (D) adults.

### Behavior of Published Signatures Over Five Seasons

To link our findings to previously identified influenza vaccine signatures, we performed a comprehensive assessment of the behavior of 1,603 previously published individual gene and gene module signatures in our data set. We manually curated published signatures from studies that carried out transcriptional profiling on adult cohorts after influenza vaccination (9, 11, 13, 15, 27, 53). We further limited the signatures to shared time points post-vaccination. This set of findings describe 935 response-associated and 653 temporal signatures in B cells and PBMCs as well as 15 age-associated signatures (*SI File 10*).

Most of the previously published signatures we validated in our data were single genes induced 7 days post-vaccination in PBMCs or B cells (*SI Appendix*, Fig. S9). Of the 135 signatures that showed significant differential expression (p < 0.001), 103 changed in the same direction as the published signature. In PBMCs we validated 26 D7 vaccine-induced genes including four genes independently discovered in our meta-analysis: *CD38, ITM2C, TNFRSF17*, and *SPATS2* (*SI Appendix*, Fig. S9B) (11). *CD38* is upregulated on the surface of antibody secreting cells, and *TNFRSF17*, or B cell maturation antigen (*BCMA*) is a receptor for B cell activating factor (*BAFF*) expressed on memory B cells and plasma cells (54). Notably, validated vaccine-induced genes in B cells include several associated with mitochondrial function whose expression was upregulated at Day 7, including *UQCRQ* (ubiquinol cytochrome c reductase, complex III, subunit VII), *ME2* (NAD-dependent malic enzyme), *TAL* (transaldolase 1), and *GLDC* (glycine decarboxylase) (*SI Appendix*, Fig. S9A). We validated several modules significantly associated with antibody response at baseline in young and older adults (*SI Appendix*, Fig. S9D) (13). Of these modules, one positively associated with antibody response (*enriched in B cells (I) (M47.0*)) is enriched in our baseline predictive model of young adults and three negatively associated with antibody response are enriched in our baseline predictive model of older adults (*Monocyte surface signature (S4), myeloid cell enriched receptors and transporters (M4.3), enriched in monocytes (II) (M11.0)*). Interestingly, these latter three modules are also enriched in predictive models of HR vs LR from D7 fold changes. Finally, there are seven validated single genes whose fold change at D7 is positively associated with antibody response in young adults (*SI Appendix*, Fig. S9C) (11). One of these genes, *HSP90B1*, or *gp96* – an ER-based chaperone protein implicated in innate and adaptive immune function – is also selected as a predictive gene of antibody response (55, 56).

## Discussion

This study is the first to evaluate the transcriptomic response to influenza vaccination in young and older adults in five consecutive vaccine seasons with three different vaccine compositions. We sought to address whether common signatures of vaccine response or transcriptional predictors of antibody response could be elucidated despite differences in seasonal vaccine composition.

To adjust for the inverse relationship between baseline antibody titers and vaccine-induced antibody production, we developed a novel vaccine response endpoint, maxRBA, to automatically correct for variation in baseline titers; this allowed us to demonstrate an age-associated decrease in antibody response in gender-matched participants. Comparing the transcriptional profiles across five seasons revealed substantial seasonal variability in both the magnitude as well as direction of response. For example, the vaccines administered in the 2010 and 2011 seasons elicited large changes in gene expression, but no statistically significant DEGs were found in the 2014 season despite a comparable sample size. Potentially, the large transcriptional changes observed in 2010 and 2011 could reflect the introduction of the A/California/7/2009 viral pandemic strain to the seasonal vaccine (as well as a change in the H3N2 vaccine strain beginning in 2010—the only year of the five studied when both influenza A strains changed). Notably, a principal component analysis revealed similar vaccine-induced signatures in the 2011 and 2014 seasons and in the 2012 and 2013 seasons. The similarities between the 2011 and 2014 seasons are intriguing because in both seasons the composition of the vaccine was identical to that in the preceding year, perhaps suggesting that these gene signatures reflect a relatively recent recall response. In contrast, the 2012 and 2013 vaccines each contained two strains which had not been present in the previous year’s vaccine. We did not observe the same trends in older adults; nonetheless, our results indicate that changes in vaccine composition, influencing factors such as vaccine strain immunogenicity and the effects of previous vaccination or infection, can alter the transcriptional response to influenza immunization.

Despite substantial inter-season variability, we identified shared vaccine-induced signatures in both young and older adults at D28. We expected D28 expression profiles to be similar to baseline; however, there were numerous transcriptional changes at D28 that were consistent across seasons with different vaccine compositions. Some of the most significant changes identified from single-season differential expression analysis in four out of five seasons were in *DUSP1, DUSP2*, and *CCL3L3*; moreover, *DUSP2* expression was also decreased in sorted CD4+ and CD8+ T cells from both young and older adults at D28. It is notable that a basal age-related alteration in phosphorylation of *DUSP1*, a negative regulator of IL-10 production, was associated with increased expression of IL-10 in monocytes from older adults (seen pre- and post-influenza vaccination) (57) and that increased *DUSP6* expression was associated with impaired T cell receptor signaling in CD4+ T cells from older adults (58). These results emphasize the importance of modulation of MAP kinase function, such as through phosphatases of the DUSP family, in the regulation of influenza vaccine response. Surprisingly, early response signatures at D2 and D7 post-vaccination were not as consistent across seasons as D28 signatures in a meta-analysis of genes and gene modules. One potential hypothesis that explains this observation is that temporal variations in early responses across seasons were not captured at the time points used, and that responses at D28 are less variable, and thus were captured in every season. It is possible that this common transcriptional program at D28 reflects a convergence towards resolution of the vaccine response in both young and older adults. However, a substantial number of BTMs showed upregulated activity at D28 without evidence of resolution to baseline, particularly in young adults; notably, we previously found evidence of enhanced TNF-alpha and IL-6 production in monocytes 28 days post-influenza immunization (57) that was blunted in monocytes from older adults. Thus, it remains possible that the transcriptional signature we observed also reflects elements of an ongoing immune activated state several weeks after vaccination.

We built predictive models of antibody response from post-vaccination transcriptional responses which were successfully validated in an independent cohort of young adults. Although transcriptional changes were correlated between age groups at D28, models of antibody response built in young adults did not validate in older adults. Strikingly, we identified a genome-wide inverse correlation between the effect size of genes discriminating HR and LR at D28 and confirmed this finding with both HAI and VNA titers. A similar inverse correlation related to age was recently reported using baseline (D0) gene expression signatures (15). We identified a single gene, *KLRB1*, whose expression alone predicted response in both age groups but in opposite directions. In young adults, changes in *KLRB1* expression were also observed in sorted CD4 and CD8 T cells, perhaps reflecting the finding that *KLRB1* expression is increased in populations of memory T cells (59). Furthermore, *KLRB1^hi^* CD8+ T cells are self-renewing memory cells that are able to reconstitute the memory T cell pool after chemotherapy (60). Thus, *KLRB1* induction in young adults may reflect an increase in memory T cell populations. In older adults, these expression patterns were not observed in sorted T cells, implying that *KLRB1* expression in another cell type, perhaps NK cells or Th17 cells, was the basis for the predictive performance.

We also built and validated predictive models of antibody response in young and older adults from D0 gene expression data. One of the predictive genes in young adults, *VASH1*, showed evidence of genetic regulation of gene expression in a previous study of influenza vaccination, suggesting that genotype may have predictive power to explain the antibody response (8). Leukocyte migration and a B cell surface signature were enriched in the predictive models. This is consistent with a recently reported meta-analysis which included baseline transcriptional profiles from the 2010, 2011, and 2012 seasons of the present study and validated a temporally stable B cell receptor signaling gene module that positively predicted response at baseline (15). While the *B cell surface signature (S2)* module we identified was not the same one identified in the previous study, our findings further support the implication of B cell transcriptional signatures as pre-vaccine biomarkers of antibody response in young adults. In older adults, we incorporated prior knowledge on gene coexpression using LogMiNeR to identify monocyte signatures which were enriched in the predictive models and were negatively associated with antibody response. Our model validated on one older adult cohort (9) but not another (13); this may reflect substantial variability in cohorts of older adults, which would be expected to be more heterogeneous in terms of comorbid medical conditions or medication use compared to young adults. Finally, we linked our findings to previously identified influenza vaccination signatures by performing a comprehensive assessment of 1,603 previously published individual gene and gene module signatures. We present the signatures that validate in any season or a meta-analysis of all seasons of our data to highlight the most consistent set of genes and gene modules associated with vaccination or antibody response in PBMC and B cells.

In summary, we profiled nearly 300 young and older adults across five vaccination seasons and, despite substantial seasonal variability in vaccine-induced transcriptional signatures, identified a core transcriptional signature shared between seasons and across age groups 28 days post-vaccination. In addition, we defined a new endpoint (maxRBA) to capture antibody response relative to baseline titer and were able to predict response in young and older adults separately using baseline transcriptional profiles. Our results suggest that vaccine composition, in concert with differences in pre-existing immunity and other individual factors, dramatically influences immune response to inactivated influenza vaccination. Furthermore, this work is a step toward understanding the underlying mechanisms of response in older adults which may be beneficial for rationally designing more effective vaccines.

## Acknowledgments

We gratefully acknowledge Dr. Randy Albrecht and Dr. Adolfo Garcia-Sastre at the Icahn School of Medicine at Mount Sinai, who led the Human Immunology Project Consortium (HIPC) core for influenza viral neutralization assays. This work was supported by NIH grant U19 AI089992, K24 AG042489, and by the Claude D. Pepper Older Americans Independence Center at Yale (to H.J.Z. and A.C.S.: P30 AG021342). Computational resources and support were provided by the Yale Center for Research Computing [NIH grants RR19895 and RR029676-01]. H.J.Z. was supported by a GEMSSTAR award from NIA (R03 AG050947). D.G.C. was supported by NIH training grant T32 EB019941. S.A. was supported by the NSF Graduate Research Fellowship Program [grant number DGE-1122492]. Any opinions, findings and conclusions or recommendations expressed in this material are those of the authors and do not necessarily reflect the views of the National Science Foundation.

## Author Contributions

Conceptualization, S.M.K., A.C.S., and S.H.K.; Software, S.A.; Formal Analysis, S.A., D.G.C., and H.M.; Investigation, S.M., H.J.Z., T.B., I.U., K.P., T.P.B, and R.B.B.; Data Curation, S.T. and H.M.; Writing – Original Draft, S.A., A.C.S., and S.H.K.; Writing – Review & Editing, All Authors; Visualization, S.A., D.G.C.

## References

1. Ohmit, S. E., J. C. Victor, E. R. Teich, R. K. Truscon, J. R. Rotthoff, D. W. Newton, S. a Campbell, M. L. Boulton, and A. S. Monto. 2008. Prevention of symptomatic seasonal influenza in 2005-2006 by inactivated and live attenuated vaccines. J. Infect. Dis. 198: 312–317.

2. Frey, S., T. Vesikari, A. Szymczakiewicz-Multanowska, M. Lattanzi, A. Izu, N. Groth, and S. Holmes. 2010. Clinical Efficacy of Cell Culture-Derived and Egg-Derived Inactivated Subunit Influenza Vaccines in Healthy Adults. Clin. Infect. Dis. 51: 997–1004.

3. Jackson, L. a, M. J. Gaglani, H. L. Keyserling, J. Balser, N. Bouveret, L. Fries, and J. J. Treanor. 2010. Safety, efficacy, and immunogenicity of an inactivated influenza vaccine in healthy adults: a randomized, placebo-controlled trial over two influenza seasons. BMC Infect. Dis. 10: 71.

4. Monto, A. S., S. E. Ohmit, J. G. Petrie, E. Johnson, R. Truscon, E. Teich, J. Rotthoff, M. Boulton, and J. C. Victor. 2009. Comparative efficacy of inactivated and live attenuated influenza vaccines. N. Engl. J. Med. 361: 1260–1267.

5. Beran, J., V. Wertzova, K. Honegr, E. Kaliskova, M. Havlickova, J. Havlik, H. Jirincova, P. Van Belle, V. Jain, B. Innis, and J.-M. Devaster. 2009. Challenge of conducting a placebo-controlled randomized efficacy study for influenza vaccine in a season with low attack rate and a mismatched vaccine B strain: a concrete example. BMC Infect. Dis. 9: 2.

6. Goodwin, K., C. Viboud, and L. Simonsen. 2006. Antibody response to influenza vaccination in the elderly: A quantitative review. Vaccine 24: 1159–1169.

7. Bucasas, K. L., L. M. Franco, C. a Shaw, M. S. Bray, J. M. Wells, D. Niño, N. Arden, J. M. Quarles, R. B. Couch, and J. W. Belmont. 2011. Early patterns of gene expression correlate with the humoral immune response to influenza vaccination in humans. J. Infect. Dis. 203: 921–9.

8. Franco, L. M., K. L. Bucasas, J. M. Wells, D. Niño, X. Wang, G. E. Zapata, N. Arden, A. Renwick, P. Yu, J. M. Quarles, M. S. Bray, R. B. Couch, J. W. Belmont, and C. a Shaw. 2013. Integrative genomic analysis of the human immune response to influenza vaccination. Elife 2: e00299.

9. Furman, D., V. Jojic, B. Kidd, S. Shen-Orr, J. Price, J. Jarrell, T. Tse, H. Huang, P. Lund, H. T. Maecker, P. J. Utz, C. L. Dekker, D. Koller, and M. M. Davis. 2013. Apoptosis and other immune biomarkers predict influenza vaccine responsiveness. Mol. Syst. Biol. 9: 659.

10. Tan, Y., P. Tamayo, H. Nakaya, B. Pulendran, J. P. Mesirov, and W. N. Haining. 2014. Gene signatures related to B-cell proliferation predict influenza vaccine-induced antibody response. Eur. J. Immunol. 44: 285–295.

11. Tsang, J. S., P. L. Schwartzberg, Y. Kotliarov, A. Biancotto, Z. Xie, R. N. Germain, E. Wang, M. J. Olnes, M. Narayanan, H. Golding, S. Moir, H. B. Dickler, S. Perl, and F. Cheung. 2014. Global analyses of human immune variation reveal baseline predictors of postvaccination responses. Cell 157: 499–513.

12. Obermoser, G., S. Presnell, K. Domico, H. Xu, Y. Wang, E. Anguiano, L. Thompson-Snipes, R. Ranganathan, B. Zeitner, A. Bjork, D. Anderson, C. Speake, E. Ruchaud, J. Skinner, L. Alsina, M. Sharma, H. Dutartre, A. Cepika, E. Israelsson, P. Nguyen, Q. A. Nguyen, a. C. Harrod, S. M. Zurawski, V. Pascual, H. Ueno, G. T. Nepom, C. Quinn, D. Blankenship, K. Palucka, J. Banchereau, and D. Chaussabel. 2013. Systems scale interactive exploration reveals quantitative and qualitative differences in response to influenza and pneumococcal vaccines. Immunity 38: 831–844.

13. Nakaya, H. I., T. Hagan, S. S. Duraisingham, E. K. Lee, M. Kwissa, N. Rouphael, D. Frasca, M. Gersten, A. K. Mehta, R. Gaujoux, G. M. Li, S. Gupta, R. Ahmed, M. J. Mulligan, S. Shen-Orr, B. B. Blomberg, S. Subramaniam, and B. Pulendran. 2015. Systems Analysis of Immunity to Influenza Vaccination across Multiple Years and in Diverse Populations Reveals Shared Molecular Signatures. Immunity 43: 1186–1198.

14. Thakar, J., S. Mohanty, A. P. West, S. R. Joshi, I. Ueda, J. Wilson, H. Meng, T. P. Blevins, S. Tsang, M. Trentalange, B. Siconolfi, K. Park, T. M. Gill, R. B. Belshe, S. M. Kaech, G. S. Shadel, S. H. Kleinstein, and A. C. Shaw. 2015. Aging-dependent alterations in gene expression and a mitochondrial signature of responsiveness to human influenza vaccination. Aging (Albany. NY). 7: 38–52.

15. HIPC-CHI Signatures Project Team, T., and T. HIPC-I Consortium. 2017. Multicohort analysis reveals baseline transcriptional predictors of influenza vaccination responses. Sci. Immunol. 2: eaal4656.

16. Song, J. Y., H. J. Cheong, I. S. Hwang, W. S. Choi, Y. M. Jo, D. W. Park, G. J. Cho, T. G. Hwang, and W. J. Kim. 2010. Long-term immunogenicity of influenza vaccine among the elderly: Risk factors for poor immune response and persistence. Vaccine 28: 3929–3935.

17. Yaari, G., C. R. Bolen, J. Thakar, and S. H. Kleinstein. 2013. Quantitative set analysis for gene expression: A method to quantify gene set differential expression including gene-gene correlations. Nucleic Acids Res. 41: e170.

18. Proost, P., P. Menten, S. Struyf, E. Schutyser, I. De Meester, and J. Van Damme. 2000. Cleavage by CD26 / dipeptidyl peptidase IV converts the chemokine LD78β into a most efficient monocyte attractant and CCR1 agonist. Blood 96: 1674–1680.

19. Kwak, S. P., D. J. Hakes, K. J. Martell, and J. E. Dixon. 1994. Isolation and Characterization of a Human Dual Specificity Protein-Tyrosine Phosphatase Gene. J. Biol. Chem. 269: 3596–3604.

20. Rohan, P. J., P. Davis, C. A. Moskaluk, M. Kearns, P. J. Rohan, P. Davis, C. A. Moskaluk, M. Kearns, H. Krutzsch, U. Siebenlist, and K. Kelly. 1993. PAC-1 : A Mitogen-Induced Nuclear Protein Tyrosine Phosphatase. Science (80-.). 259: 1763–1766.

21. Wei, W., Y. Jiao, A. Postlethwaite, J. M. Stuart, Y. Wang, D. Sun, and W. Gu. 2013. Dual-specificity phosphatases 2: surprising positive effect at the molecular level and a potential biomarker of diseases. Genes Immun. 14: 1–6.

22. Mohanty, S., S. R. Joshi, I. Ueda, J. Wilson, T. P. Blevins, B. Siconolfi, H. Meng, L. Devine, K. Raddassi, S. Tsang, R. B. Belshe, D. A. Hafler, S. M. Kaech, S. H. Kleinstein, M. Trentalange, H. G. Allore, and A. C. Shaw. 2015. Prolonged proinflammatory cytokine production in monocytes modulated by interleukin 10 after influenza vaccination in older adults. J. Infect. Dis. 211: 1174–1184.

23. Li, S., N. Rouphael, S. Duraisingham, S. Romero-Steiner, S. Presnell, C. Davis, D. S. Schmidt, S. E. Johnson, A. Milton, G. Rajam, S. Kasturi, G. M. Carlone, C. Quinn, D. Chaussabel, a K. Palucka, M. J. Mulligan, R. Ahmed, D. S. Stephens, H. I. Nakaya, and B. Pulendran. 2014. Molecular signatures of antibody responses derived from a systems biology study of five human vaccines. Nat. Immunol. 15: 195–204.

24. Kanehisa, M., and S. Goto. 2000. KEGG: Kyoto encyclopedia of genes and genomes. Nucleic Acids Res. 28: 27–30.

25. Abbas, a R., D. Baldwin, Y. Ma, W. Ouyang, A. Gurney, F. Martin, S. Fong, M. van Lookeren Campagne, P. Godowski, P. M. Williams, a C. Chan, and H. F. Clark. 2005. Immune response in silico (IRIS): immune-specific genes identified from a compendium of microarray expression data. Genes Immun. 6: 319–331.

26. Avey, S., S. Mohanty, J. Wilson, H. Zapata, S. R. Joshi, B. Siconolfi, S. Tsang, A. C. Shaw, and S. H. Kleinstein. 2017. Multiple network-constrained regressions expand insights into influenza vaccination responses. Bioinformatics 33: i208–i216.

27. Nakaya, H. I., J. Wrammert, E. K. Lee, L. Racioppi, S. Marie-Kunze, W. N. Haining, A. R. Means, S. P. Kasturi, N. Khan, G.-M. Li, M. McCausland, V. Kanchan, K. E. Kokko, S. Li, R. Elbein, A. K. Mehta, A. Aderem, K. Subbarao, R. Ahmed, and B. Pulendran. 2011. Systems biology of vaccination for seasonal influenza in humans. Nat. Immunol. 12: 786–795.

28. Gaucher, D., R. Therrien, N. Kettaf, B. R. Angermann, G. Boucher, A. Filali-Mouhim, J. M. Moser, R. S. Mehta, D. R. Drake, E. Castro, R. Akondy, A. Rinfret, B. Yassine-Diab, E. a Said, Y. Chouikh, M. J. Cameron, R. Clum, D. Kelvin, R. Somogyi, L. D. Greller, R. S. Balderas, P. Wilkinson, G. Pantaleo, J. Tartaglia, E. K. Haddad, and R.-P. Sékaly. 2008. Yellow fever vaccine induces integrated multilineage and polyfunctional immune responses. J. Exp. Med. 205: 3119–3131.

29. Querec, T. D., R. S. Akondy, E. K. Lee, W. Cao, H. I. Nakaya, D. Teuwen, A. Pirani, K. Gernert, J. Deng, B. Marzolf, K. Kennedy, H. Wu, S. Bennouna, H. Oluoch, J. Miller, R. Z. Vencio, M. Mulligan, A. Aderem, R. Ahmed, and B. Pulendran. 2009. Systems biology approach predicts immunogenicity of the yellow fever vaccine in humans. Nat. Immunol. 10: 116–125.

30. Mitchell, P., and D. Tollervey. 2000. mRNA stability in eukaryotes. Curr. Opin. Genet. Dev. 10: 193–198.

31. Molleston, J. M., and S. Cherry. 2017. Attacked from all sides: RNA decay in antiviral defense. Viruses 9.

32. Liu, S. W., G. C. Katsafanas, R. Liu, L. S. Wyatt, and B. Moss. 2015. Poxvirus decapping enzymes enhance virulence by preventing the accumulation of dsRNA and the induction of innate antiviral responses. Cell Host Microbe 17: 320–331.

33. Khaperskyy, D. A., S. Schmaling, J. Larkins-Ford, C. McCormick, and M. M. Gaglia. 2016. Selective Degradation of Host RNA Polymerase II Transcripts by Influenza A Virus PA-X Host Shutoff Protein. PLoS Pathog. 12: 1–25.

34. Gaglia, M. M., S. Covarrubias, W. Wong, and B. A. Glaunsinger. 2012. A common strategy for host RNA degradation by divergent viruses. J Virol 86: 9527–9530.

35. Patwari, P., and R. T. Lee. 2012. An expanded family of arrestins regulate metabolism. Trends Endocrinol. Metab. 23: 216–222.

36. Nakamura, N., and S. Hirose. 2008. Regulation of Mitochondrial Morphology by USP30, a Deubiquitinating Enzyme Present in the Mitochondrial Outer Membrane. Mol. Biol. Cell 19: 1903–1911.

37. Twyffels, L., C. Gueydan, and V. Kruys. 2014. Transportin-1 and Transportin-2: Protein nuclear import and beyond. FEBS Lett. 588: 1857–1868.

38. Meng, H., G. Yaari, C. R. Bolen, S. Avey, and S. H. Kleinstein. 2019. Gene set meta-analysis with Quantitative Set Analysis for Gene Expression (QuSAGE). PLOS Comput. Biol. 15: e1006899.

39. Curran, J. E., J. B. M. Jowett, K. S. Elliott, Y. Gao, K. Gluschenko, J. Wang, D. M. Abel Azim, G. Cai, M. C. Mahaney, A. G. Comuzzie, T. D. Dyer, K. R. Walder, P. Zimmet, J. W. MacCluer, G. R. Collier, A. H. Kissebah, and J. Blangero. 2005. Genetic variation in selenoprotein S influences inflammatory response. Nat. Genet. 37: 1234–1241.

40. Ye, Y., Y. Shibata, C. Yun, D. Ron, and T. A. Rapoport. 2004. A membrane protein complex mediates retro-translocation from the ER lumen into the cytosol. Nature 429: 841–847.

41. Jeffrey, K. L., M. Camps, C. Rommel, and C. R. Mackay. 2007. Targeting dual-specificity phosphatases: manipulating MAP kinase signalling and immune responses. Nat. Rev. Drug Discov. 6: 391–403.

42. Kutty, R. G., G. Xin, D. M. Schauder, S. M. Cossette, M. Bordas, W. Cui, and R. Ramchandran. 2016. Dual specificity phosphatase 5 is essential for T cell survival. PLoS One 11: 1–16.

43. Indiveri, C., V. Iacobazzi, A. Tonazzi, N. Giangregorio, V. Infantino, P. Convertini, L. Console, and F. Palmieri. 2011. The mitochondrial carnitine/acylcarnitine carrier: Function, structure and physiopathology. Mol. Aspects Med. 32: 223–233.

44. van Duin, D., H. G. Allore, S. Mohanty, S. Ginter, F. K. Newman, R. B. Belshe, R. Medzhitov, and A. C. Shaw. 2007. Prevaccine Determination of the Expression of Costimulatory B7 Molecules in Activated Monocytes Predicts Influenza Vaccine Responses in Young and Older Adults. J. Infect. Dis. 195: 1590–1597.

45. Panda, A., F. Qian, S. Mohanty, D. van Duin, F. K. Newman, L. Zhang, S. Chen, V. Towle, R. B. Belshe, E. Fikrig, H. G. Allore, R. R. Montgomery, and A. C. Shaw. 2010. Age-Associated Decrease in TLR Function in Primary Human Dendritic Cells Predicts Influenza Vaccine Response. J. Immunol. 184: 2518–2527.

46. Aldemir, H., V. Prod’homme, M.-J. Dumaurier, C. Retiere, G. Poupon, J. Cazareth, F. Bihl, and V. M. Braud. 2005. Cutting Edge: Lectin-Like Transcript 1 Is a Ligand for the CD161 Receptor. J. Immunol. 175: 7791–7795.

47. Rosen, D. B., J. Bettadapura, M. Alsharifi, P. A. Mathew, H. S. Warren, and L. L. Lanier. 2005. Cutting Edge: Lectin-Like Transcript-1 Is a Ligand for the Inhibitory Human NKR-P1A Receptor. J. Immunol. 175: 7796–7799.

48. Kleinschek, M. A., K. Boniface, S. Sadekova, J. Grein, E. E. Murphy, S. P. Turner, L. Raskin, B. Desai, W. A. Faubion, R. de Waal Malefyt, R. H. Pierce, T. McClanahan, and R. A. Kastelein. 2009. Circulating and gut-resident human Th17 cells express CD161 and promote intestinal inflammation. J. Exp. Med. 206: 525–534.

49. Maggi, L., V. Santarlasci, M. Capone, A. Peired, F. Frosali, S. Q. Crome, V. Querci, M. Fambrini, F. Liotta, M. K. Levings, E. Maggi, L. Cosmi, S. Romagnani, and F. Annunziato. 2010. CD161 is a marker of all human IL-17-producing T-cell subsets and is induced by RORC. Eur. J. Immunol. 40: 2174–2181.

50. Cosmi, L., R. De Palma, V. Santarlasci, L. Maggi, M. Capone, F. Frosali, G. Rodolico, V. Querci, G. Abbate, R. Angeli, L. Berrino, M. Fambrini, M. Caproni, F. Tonelli, E. Lazzeri, P. Parronchi, F. Liotta, E. Maggi, S. Romagnani, and F. Annunziato. 2008. Human interleukin 17–producing cells originate from a CD161 ^+^ CD4 ^+^ T cell precursor. J. Exp. Med. 205: 1903–1916.

51. Gorenshteyn, D., E. Zaslavsky, M. Fribourg, C. Y. Park, A. K. Wong, A. Tadych, B. M. Hartmann, R. A. Albrecht, A. García-Sastre, S. H. Kleinstein, O. G. Troyanskaya, and S. C. Sealfon. 2015. Interactive Big Data Resource to Elucidate Human Immune Pathways and Diseases. Immunity 43: 605–614.

52. Cho, N. E., B. R. Bang, P. Gurung, M. Li, D. L. Clemens, T. M. Underhill, L. P. James, J. R. Chase, and T. Saito. 2016. Retinoid regulation of antiviral innate immunity in hepatocytes. Hepatology 63: 1783–1795.

53. Furman, D., J. Chang, L. Lartigue, C. R. Bolen, F. Haddad, B. Gaudilliere, E. A. Ganio, G. K. Fragiadakis, M. H. Spitzer, I. Douchet, S. Daburon, J.-F. Moreau, G. P. Nolan, P. Blanco, J. Déchanet-Merville, C. L. Dekker, V. Jojic, C. J. Kuo, M. M. Davis, and B. Faustin. 2017. Expression of specific inflammasome gene modules stratifies older individuals into two extreme clinical and immunological states. Nat. Med..

54. Darce, J. R., B. K. Arendt, X. Wu, and D. F. Jelinek. 2014. Regulated Expression of BAFF-Binding Receptors during Human B Cell Differentiation. J. Immunol. 179: 7276–7286.

55. Pockley, A. G., and B. Henderson. 2018. Extracellular cell stress (Heat shock) proteins—immune responses and disease: An overview. Philos. Trans. R. Soc. B Biol. Sci. 373.

56. Randow, F., and B. Seed. 2001. Endoplasmic reticulum chaperone gp96 is required for innate immunity but not cell viability. Nat. Cell Biol. 3: 891–896.

57. Mohanty, S., S. R. Joshi, I. Ueda, J. Wilson, T. P. Blevins, B. Siconolfi, H. Meng, L. Devine, K. Raddassi, S. Tsang, R. B. Belshe, D. a. Hafler, S. M. Kaech, S. H. Kleinstein, M. Trentalange, H. G. Allore, and a. C. Shaw. 2014. Prolonged Proinflammatory Cytokine Production in Monocytes Modulated by Interleukin 10 After Influenza Vaccination in Older Adults. J. Infect. Dis. 211: 1174–1184.

58. Li, G., M. Yu, W. W. Lee, M. Tsang, E. Krishnan, C. M. Weyand, and J. J. Goronzy. 2012. Decline in miR-181a expression with age impairs T cell receptor sensitivity by increasing DUSP6 activity. Nat. Med. 18: 1518–1524.

59. Kirkham, C. L., and J. R. Carlyle. 2014. Complexity and Diversity of the NKR-P1:Clr (Klrb1:Clec2) Recognition Systems. Front. Immunol. 5: 1–16.

60. Turtle, C. J., H. M. Swanson, N. Fujii, E. H. Estey, and S. R. Riddell. 2009. A Distinct Subset of Self-Renewing Human Memory CD8+ T Cells Survives Cytotoxic Chemotherapy. Immunity 31: 834–844.

