## Supplementary material for "Seasonal Variability and Shared Molecular Signatures of Inactivated Influenza Vaccination in Young and Older Adults": SI Appendix

[a] Interdepartmental Program in Computational Biology and Bioinformatics, Yale University, New Haven, CT 06511, USA
[b] Section of Infectious Diseases, Department of Internal Medicine, Yale School of Medicine, New Haven, CT 06520, USA
[c] Department of Pathology, Yale School of Medicine, New Haven, CT 06520, USA
[d] Department of Immunobiology, Yale School of Medicine, New Haven, CT 06520, USA
[e] Division of Infectious Diseases, Department of Medicine, Saint Louis University School of Medicine, St. Louis, MO 63104 USA
[1] S.A. and S.M. contributed equally to this work
[2] Current address: Salk Institute for Biological Studies, La Jolla, CA 92037
[3] A.C.S. and S.H.K contributed equally to this work

**This PDF file includes:**

Supplementary Information File Descriptions
Supplementary Information Text
Figs. S1 to S9

**Supplementary Information File Descriptions**

**Supplementary File 1.** Table of differentially expressed genes in each individual season.
**Supplementary File 2.** Genes and enrichments of 7 gene clusters in Fig. 2.
**Supplementary File 3.** Gene set enrichment of the first two principal components in Fig. 2.
**Supplementary File 4.** Gene module activity and p-values for each season and meta-analysis.
**Supplementary File 5.** Single gene meta-analysis results for vaccine-induced genes.
**Supplementary File 6.** Single gene meta-analysis results for young vs. older adults.
**Supplementary File 7.** Gene module meta-analysis results for young vs. older adults.
**Supplementary File 8.** Gene set enrichment for predictive models of antibody response.
**Supplementary File 9.** Single gene meta-analysis results for HR vs. LR.
**Supplementary File 10.** Validation of published influenza vaccination signatures.

### Supplementary Information Text

***Peripheral Blood Mononuclear Cell (PBMC) Isolation***

Blood samples were collected in sodium heparin tubes (BD Biosciences) from volunteers with prior consent to an IRB approved protocol. About 20 ml of fresh blood were layered on top of 20 ml Histopaque-1077 solution (Sigma-Aldrich). Samples were carefully transferred to a centrifuge (Legend XT, Thermo Fisher Scientific) and centrifuged for 20 minutes at room temperature without brake at 2200 rpm. Buffy coats containing peripheral blood mononuclear cells (PBMC) at the interface were carefully collected to 50 ml RPMI medium with 10% BSA. Samples were centrifuged to pellet cells and the supernatant were discarded. Lymphocyte pellets were re-suspended in 50 ml RPMI containing 10% FBS. A 20-microliter volume of suspended PMBC was mixed with equal volume Trypan Blue solution (Thermo Fisher Scientific) and incubated at room temperature for about 2 minutes. A 20-microliter cell suspension was immediately counted using Haemocytometer under light microscope. Percent viability was determined by the formula (Number of total cells counted – Number of Blue cells counted) x 100. Samples with 95% or more viability were selected for all experiments including RNA extraction.

***PBMC RNA Extraction***

About 10 million cells (PBMC) in 1 ml RPMI medium from samples with 95% or more viability were taken soon after Histopaque-1077 density gradient separation for RNA extraction. PBMC in RPMI medium were centrifuged for about 10 minutes on a bench top centrifuge at 10,000 rpm. Supernatant was removed carefully from each sample to ensure a clear pellet of PBMC without residual RPMI medium. To each pellet about 700 microliters of QIAzol lysis reagent (Qiagen) was added and mixed by pipetting at least 10 times to ensure proper cell lysis. Lysed cells were immediately frozen at $-80^{0}$ C until further extraction using QIAcube instrument (Qiagen).

***QIAcube RNA Extraction Protocol***

All RNA samples were extracted using miRNeasy kit (cat. 217004, Qiagen) following the instructions provided using a QIAcube (Qiagen). Briefly, samples lysed in QIAzol reagent were incubated for 5 minutes at room temperature ($15^{0}$ C - $25^{0}$ C). To each sample about 140 microliters of chloroform was added and shaken vigorously and left at room temperature for about 2-3 minutes. Subsequently, samples were centrifuged at $4^{0}$ C at 12,000 x g for 15 minutes. Upper aqueous phase containing the RNA species were carefully transferred to a 2 ml collection tube (cat. 990381) Qiagen without touching the interphase and placed in the QIAcube for extraction. For every sample a rotor adapter was prepared with a RNeasy mini spin column at position L1 and a 1.5 ml collection tube at position L3 and placed in the QIAcube rotor. All reagents were prepared by adding proper amount of 100% ethanol (44 ml to buffer RPE and 30 ml to buffer RWT) prior to extraction and placed in respective positions in reagent rack in QIAcube (100% ethanol in position 3, Buffer RWT in position 4, buffer RPE in position 5 and RNase-free water in position 6). RNA extraction was carried out by executing recommended protocol (FIW-50-001-J_FW_MB and PLC program version FIW-50-002-G_PLC_MB) available from the QIAcube web portal. RNA samples with RIN value above 7.0 were used for gene expression analysis.

***RNA Isolation and Cell Sorting***

Peripheral Blood Mononuclear Cells (PBMCs) isolation and RNA preparation on freshly isolated PBMCs was performed as described (1). For cell sorting, frozen PBMCs were thawed, washed, and stained using antibodies directed against: CD19(HIB19), CD4(SK3), CD20(2H7), CD8(SK1), and CD3(UCHT1) (from BD Biosciences) and using a BD Fortessa instrument. T and B cells were sorted from a subgroup of samples in seasons 2010-2012. We evaluated the purity of 268 / 270 sorts for CD4+ T cells with median purity of 97%, 254 / 254 sorts for CD8+ T cells with median purity of 97%, and 252 / 256 sorts for B cells with median purity of 99%. 90% of the samples had a post-sort purity of at least 90% for all 3 cell types. Very few samples had a purity of less than 80% (0.4%, 1.1%, and 5.6% for CD4 T cells, CD8 T cells, and B cells, respectively).

***HAI and VNA Titer Measurements and Response Endpoint Definition***

Serum samples were collected pre-vaccination (D0) and 28 days post-vaccination (D28). Hemagglutination inhibition (HAI) assays were performed as previously described (2). Virus neutralization assays (VNA) were performed as described elsewhere (3, 4). Briefly, two-fold dilutions (50 µL) of the RDE-treated sera in sterile Opti-MEM (Invitrogen, Carlsbad, CA) were mixed with 200 PFU of influenza virus (5 µL). The serum-virus samples were then incubated at room temperature for 60 minutes to allow any HA-specific antibodies present in the serum to neutralize the influenza virus. The serum-virus samples (55 µL) were then transferred to MDCK cell cultures cultured in 96-wll flat bottom plates. Following virus absorption for 60 minutes, the serum-virus inocula were removed, and the MDCK cells were cultured for 4 days in Opti-MEM supplemented with 1 µg/ml of tosylsulfonyl phenylalanyl chloromethyl ketone (TPCK)-trypsin (Sigma-Aldrich). Virus production was determined by HA assay. The neutralization titer was defined as the reciprocal of the highest dilution of serum that neutralizes 200 PFU of influenza virus.

To adjust for inverse correlations between HAI titer fold changes and baseline titers, we developed an automated metric: maximum Residual after Baseline Adjustment (maxRBA). First, young and older cohorts were separated, and endpoints were calculated in each season and each age group separately. Any fold changes less than 1 were set to 1 since we did not expect the number of antibodies in the blood to decrease by 2-fold in the 28 days after vaccination and this was likely due to measurement error. Next, the baseline and fold changes were $\log_2$ transformed and an exponential curve was fit to the fold change vs. baseline titers for each strain. Then the residuals were calculated and for each subject the maximum residual across all strains was selected as the maxRBA. Finally, "high" and "low" responders were defined as the top and bottom 40th percent of maxRBA, respectively. The maxRBA values presented in Fig. 1 and Fig. S3 were calculated by combining young and older adults within each season to allow for comparison across age groups. The code to calculate maxRBA is available in the *Calculate_maxRBA()* function from the titer R package (https://bitbucket.org/kleinstein/titer).

***RNA Processing and Microarrays***

Each RNA sample was quantified, and integrity assessed by the Agilent 2100 BioAnalyser (Agilent, CA). Samples were processed for cRNA generation using the Illumina TotalPrep cRNA Amplification Kit and subsequently hybridized to the Human HT12-V4.0 BeadChip (Illumina, CA). For gene expression analyses, samples were processed and hybridized to HumanHT-12v4 Expression BeadChip (Illumina San Diego, CA). Arrays were processed at Yale's Keck Biotechnology Resource Laboratory and raw expression data were output using Illumina GenomeStudio software. Samples from each season were processed in batches and all samples from each subject were run on the same chip to mitigate batch effects. Microarray data are available through the Gene Expression Omnibus (GEO) Database with accession numbers GSE59635, GSE59654, GSE59743, GSE101709, and GSE101710 for PBMC data and GSE65440, GSE65442, and GSE95584 for B and T cell data. Data from seasons 2010 and 2011 (GSE59635 and GSE59654) were previously published (1) as well as data from D0 and D7 timepoints from the 2012 season (GSE59743) (5). The D0 expression in PBMC (GSE59635, GSE59654, GSE59743, GSE101709, and GSE101710) of a single gene, *MINCLE*, was also recently published (6). The remainder of the data in this work has not been previously published. The data were quantile normalized and $\log_2$ transformed within each season. Multiple probes were collapsed to unique Entrez Gene IDs by selecting the probe with the highest average expression. The Bioconductor package illuminaHumanv4.db version 1.26 was used to map the probes (7).

#### *Gene Module and Differential Expression Analysis*

Differentially expressed modules (DEMs) were defined at each post-vaccination time-point using QuSAGE with an FDR < 0.05 (8). Differentially expressed genes (DEGs) were defined at each post-vaccination time-point using two criteria: (1) an absolute fold-change of at least 1.25 relative to the pre-vaccination time point and (2) a significant change in expression by limma (version 3.30.7) after correction for multiple hypothesis testing (FDR < 0.05) (9) . Clusters of DEGs were determined by hierarchical clustering using Ward's minimum variance method with the distance between any 2 genes defined by 1 less the Pearson correlation coefficient. Enrichment of DEG clusters was performed using EnrichR v1.0 with the following databases: GO_Cellular_Component_2018, GO_Biological_Process_2018, GO_Molecular_Function_2018, KEGG_2016, and Reactome_2016 (10). Enrichment of principal components was performed using the geneSetTest function in the limma R package v3.24.15 (9). Control of the false discovery rate was performed according to the procedure in (11) unless otherwise noted.

#### *Meta-Analysis*

First, genes were filtered to those that were detected in at least 20% of samples with a p-value below 0.05. For the gene module meta-analysis, QuSAGE activity distributions were taken from the single season analysis and convoluted to create a probability density function for the meta-analysis (12). Modules with an FDR < 0.10 in the meta-analysis were chosen as significant. For the single-gene meta-analysis, a random-effects model was fit for every gene, at each time point, in each age group separately. This model allows for variation in effect sizes across seasons. Effect sizes (mean differences) were calculated for every gene in each season separately and then combined using the *rma()* function from the metafor R package (13). The restricted maximum-likelihood estimator for the amount of heterogeneity was used (14). Genes with an FDR < 0.05 in the meta-analysis were chosen as significant.

#### *Predicting Antibody Response from Transcriptional Profiles*

Because females tended to respond better than males, genes located on the X and Y chromosomes were removed to avoid selection of sex-linked genes that may be confounded with vaccine response. For baseline predictors, the 1,000 genes with the largest coefficient of variation were selected as the initial feature set. For post-vaccination predictors, the log fold-change from D0 was calculated for each gene, and the 1,000 genes with the largest fold-change magnitudes were selected as the initial feature set. The baseline or fold-changes were standardized by subtracting the mean and dividing by the standard deviation. Finally, this preprocessed data from each individual season was combined to form the discovery data. The young adult models were tested on GSE47353 at baseline, day 1, day 7, and day 70 while the older adult models were tested on GSE41080 at baseline and GSE74813 at baseline, day 1, day 7, and day 14 post-vaccination. The Logistic Multiple Network-constrained Regression (LogMiNeR) framework was performed as described (5). Briefly, 5-fold cross validation was used to select the optimal tuning parameters and 50 iterations of cross validation were performed on different splits of the discovery data set. The prior knowledge networks were defined for Reactome (15, 16), Gene Ontology (GO) (17), Blood Transcriptional Modules (BTM) (18) and cell-specific signatures (CELLS) (19) by connecting all pairs of genes within each gene set. The KEGG network incorporated pathway topology and was built using the KEGGgraph R package v1.26.0 (20). ImmuneGlobal (ImmuNet) and ExpOnly_ImmuneGlobal (ImmuNet_Exp) networks were obtained from ImmuNet and edges were restricted to those with confidence of at least 0.1 (21). The STRING network incorporated all experimental evidence from the STRING database v10.0 (22).

***Comparison Against Published Signatures***

Influenza vaccination signatures were manually curated from several publications (23–28). Differentially expressed genes (DEGs) were defined for each contrast using a significant change in expression by limma (version 3.30.7) ($p < 0.001$). Differentially expressed modules were defined as described above using QuSAGE but with a p-value cutoff of $p < 0.001$. Single-gene meta-analysis and gene module meta-analysis were performed as described above for each published signature without filtering lowly expressed genes. HAI-associated comparisons were discretized and tested using maxRBA-defined HR vs LR as described above.

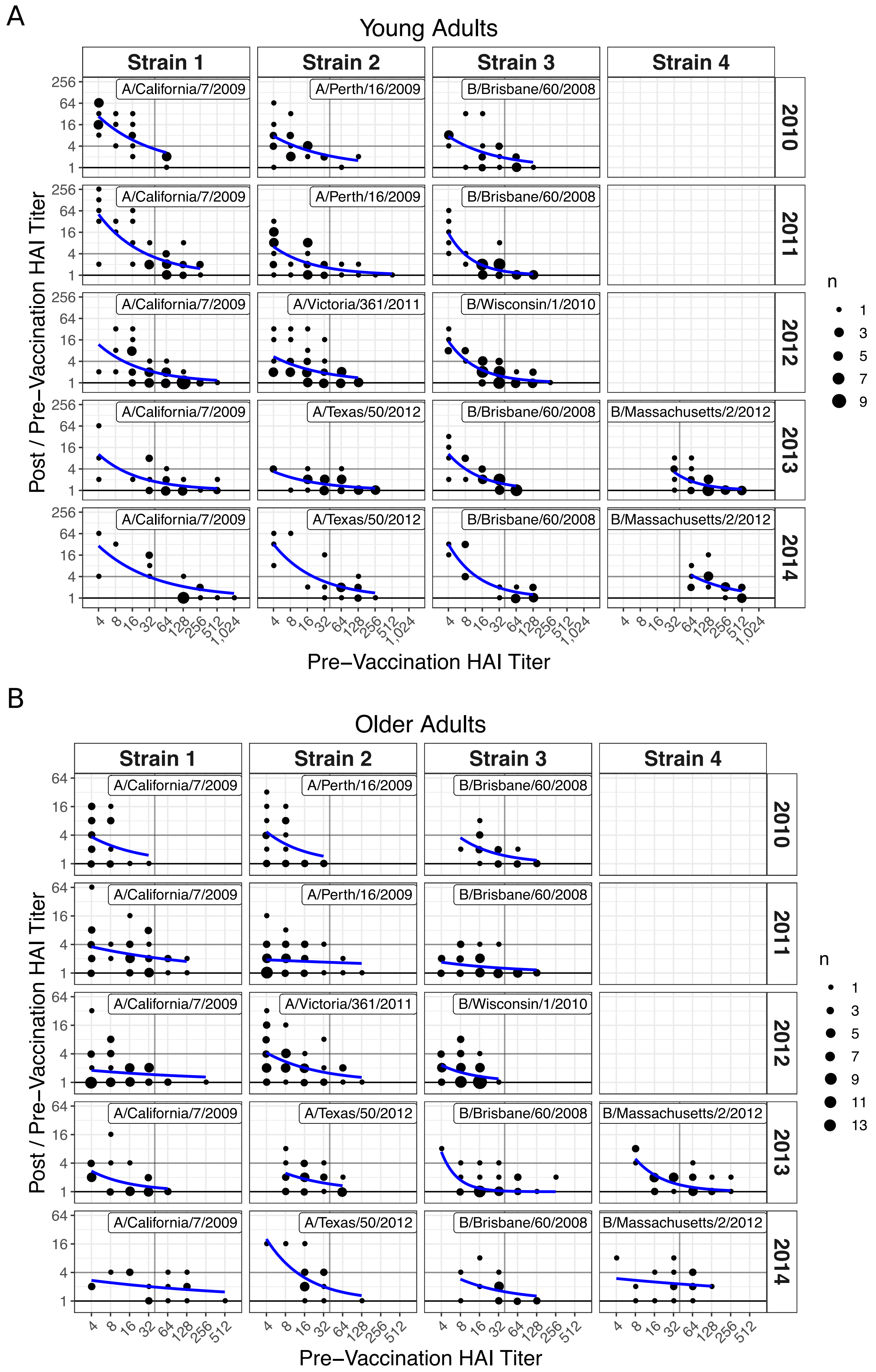
A
Young Adults
Strain 1
Strain 2
Strain 3
Strain 4
A/California/7/2009
A/Perth/16/2009
B/Brisbane/60/2008
A/Victoria/361/2011
B/Wisconsin/1/2010
A/Texas/50/2012
B/Massachusetts/2/2012
2010
2011
2012
2013
2014
Post / Pre−Vaccination HAI Titer
Pre−Vaccination HAI Titer
n
1
3
5
7
9
B
Older Adults
Strain 1
Strain 2
Strain 3
Strain 4
Post / Pre−Vaccination HAI Titer
Pre−Vaccination HAI Titer
n
1
3
5
7
9
11
13

**Fig. S1. Negative Relationship Between Baseline HAI Titer and Fold Change**

(A-B) Bubble charts of pre-vaccination HAI titer versus titer fold change post-vaccination for young (A) and older (B) adults. Plots are faceted by strain in columns and season in rows. The area of each point is proportional to the number of subjects (n) represented by each point and given in the legend. A horizontal line at fold change of 4 indicates what is typically considered a good response and a vertical line at a baseline titer of 1:40 indicates a typical cutoff for a protective titer. Blue curves show the exponential fit to the data in each facet.

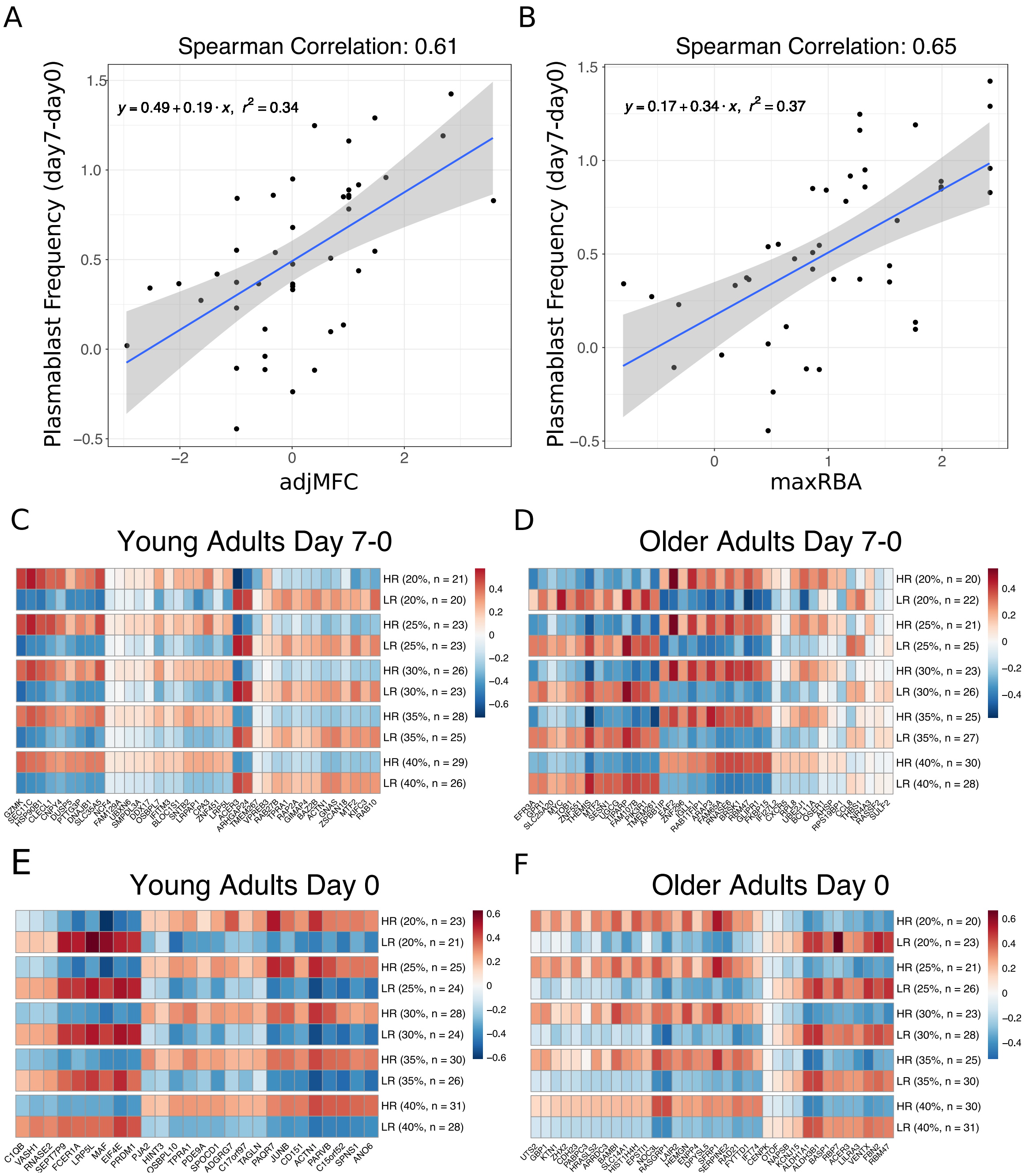


**Fig. S2. Validation of maxRBA and Sensitivity Analysis**

(A-B) Scatter plots of adjMFC (A) and maxRBA (B) versus $\log_{10}$ change in the % of CD27hi CD38hi of CD20− B cells (plasmablasts) 7 days post-vaccination in young adults from (23). (C-D) Heatmaps of average expression of predictive genes (related to Fig. 4 C-D) when varying cutoff of maxRBA from 20-50%. (E-F) Heatmaps of average expression of predictive genes (related to Fig. 5 C-D) when varying cutoff of maxRBA from 20-50%.

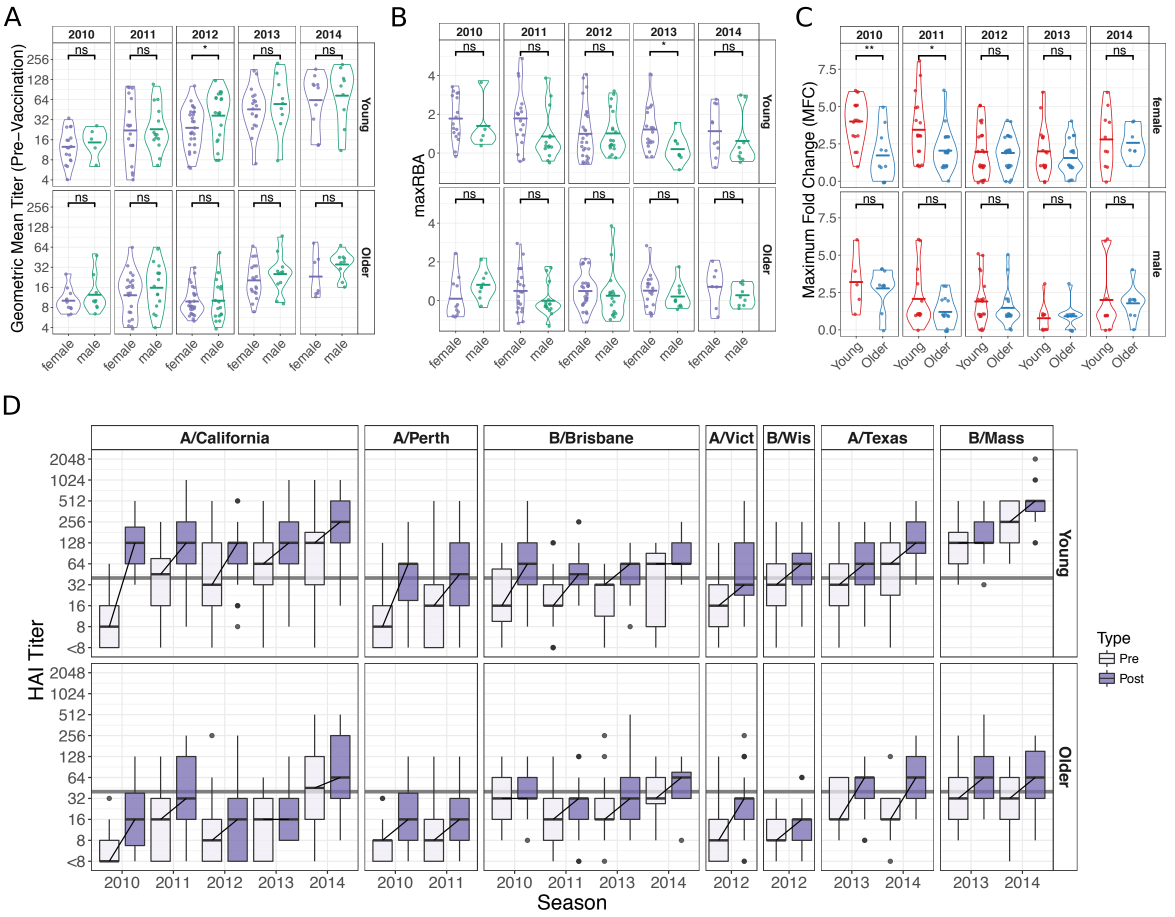


**Fig. S3. Age and Gender Differences in HAI Titers. Related to Fig. 1**

(A-B) Violin plots of pre-vaccination HAI titers (A) and HAI responses measured by maxRBA (B) are separated by season and age group to compare genders. (C) Violin plots of HAI maximum fold change separated by season and gender to compare age groups (related to Fig. 1C). Crossbars indicate the mean. Not Significant (ns) $p > 0.05$, * $p < 0.05$, ** $p < 0.01$, *** $p < 0.001$, **** $p < 0.0001$ independent two-sided Wilcoxon rank sum test. (D) Boxplots of pre- and post-vaccination HAI titers over five seasons for each individual influenza strain and age group. Lines between boxes connect the median pre-vaccination titer to the median post-vaccination titer. Strain names are abbreviated for legibility as follows: A/California = A/California/7/2009; A/Perth = A/Perth/16/2009; B/Brisbane = B/Brisbane/60/2008; A/Vict = A/Victoria/361/2011; B/Wis = B/Wisconsin/1/2010; B/Mass = B/Massachusetts/2/2012.

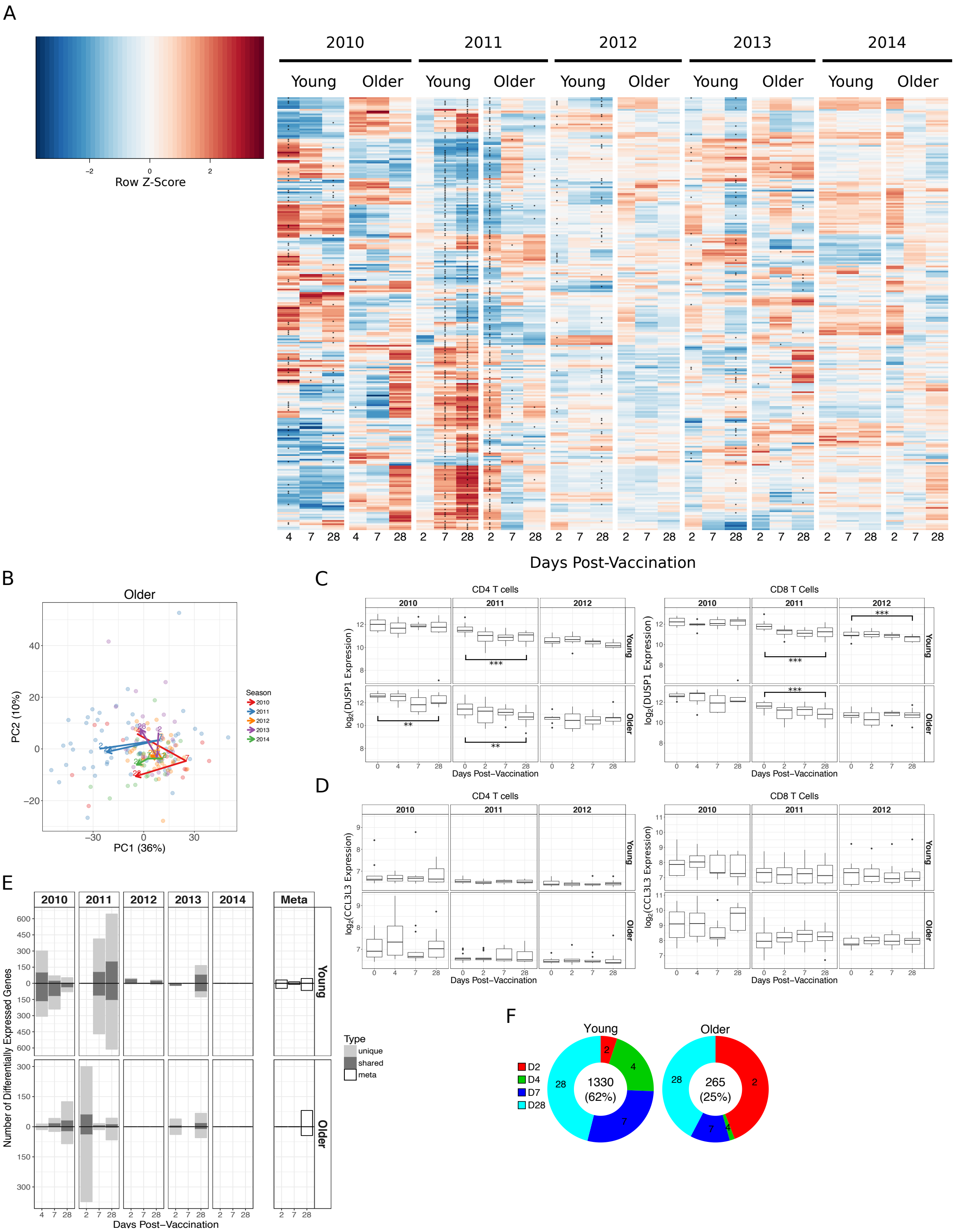
A
2010
2011
2012
2013
2014
Young
Older
Row Z-Score
Days Post-Vaccination
B
Older
PC1 (36%)
PC2 (10%)
Season
2010
2011
2012
2013
2014
C
CD4 T cells
CD8 T Cells
log2(DUSP1 Expression)
Young
Older
Days Post-Vaccination
D
CD4 T cells
CD8 T Cells
log2(CCL3L3 Expression)
Days Post-Vaccination
E
Meta
Number of Differentially Expressed Genes
Type
unique
shared
meta
Days Post-Vaccination
F
Young
Older
D2
D4
D7
D28
1330
(62%)
265
(25%)

**Fig. S4. Additional Seasonal Variability in Signatures Induced by Influenza Vaccination. Related to Fig. 2**

(A) A row-normalized heatmap of all 159 significantly differentially expressed modules. Significant modules are indicated by *. (B) The first two principal components from a principal component analysis (PCA) of all differentially expressed genes in older adults. (C and D) Expression of *DUSP1* (C) and *CCL3L3* (D) expression in sorted CD4 and CD8 T cells. ** $p < 0.01$, *** $p < 0.001$ one-sided t-test comparing day 28 and day 0 only. (E) Bar plots of the number of DEGs in each season within each age group as well as a meta-analysis of five seasons ("Meta"). (F) Donut charts of the shared genes from panel E. The size of each slice represents the fraction of shared genes from the indicated day post-vaccination while the number and fraction of shared DEGs is displayed in the center.

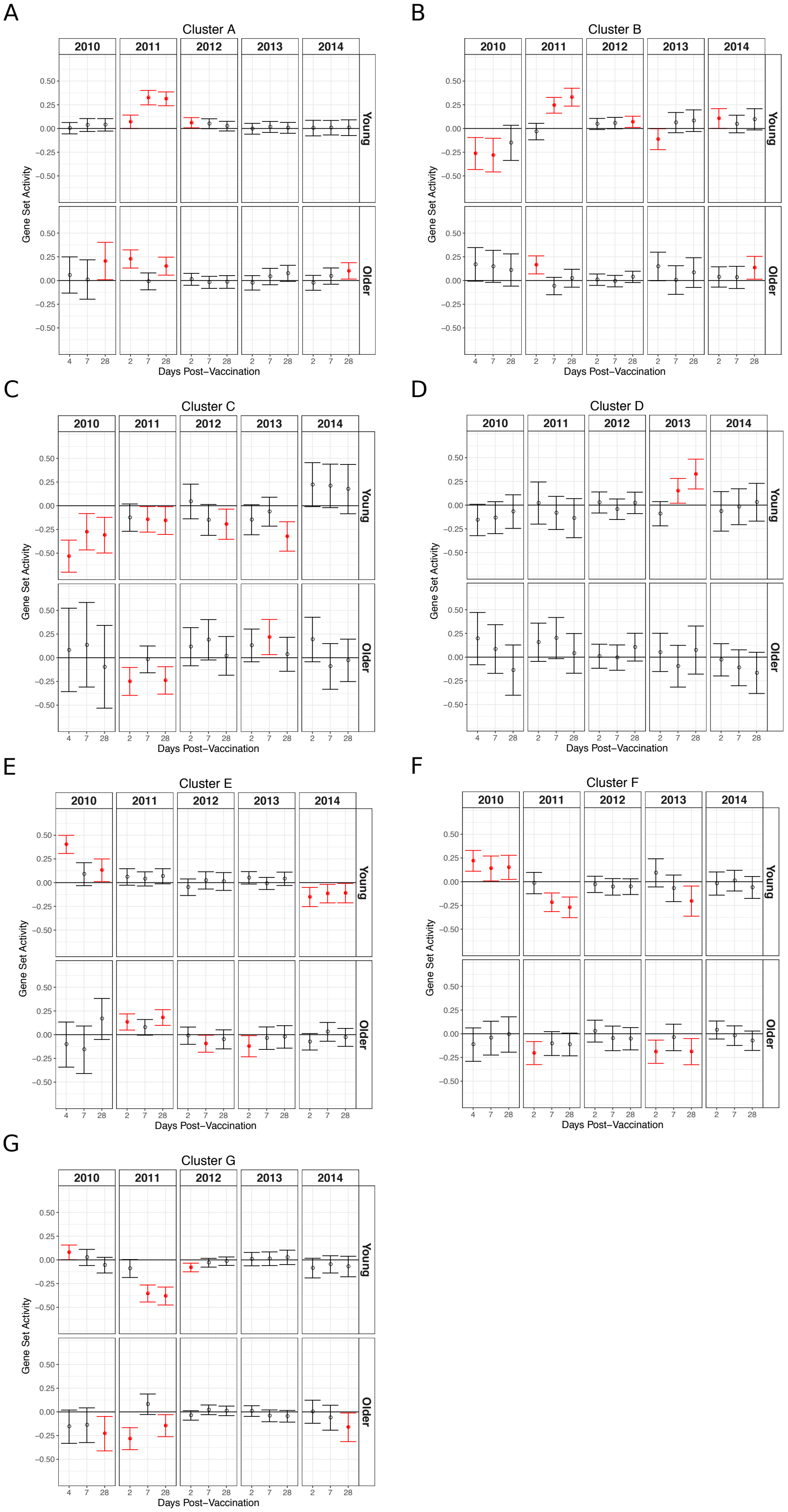
A
Cluster A
B
Cluster B
C
Cluster C
D
Cluster D
E
Cluster E
F
Cluster F
G
Cluster G
2010
2011
2012
2013
2014
Young
Older
Gene Set Activity
Days Post−Vaccination
0.50
0.25
0.00
−0.25
−0.50
4
7
28
2

**Fig. S5. Activity of Gene Clusters in Five Seasons. Related to Fig. 2**

Plots of the gene set activity calculated using QuSAGE for the 7 clusters shown in Fig. 2A. Points are means and whiskers represent the 95% confidence interval. Significant activities ($p < 0.05$) are shown in filled circles in red.

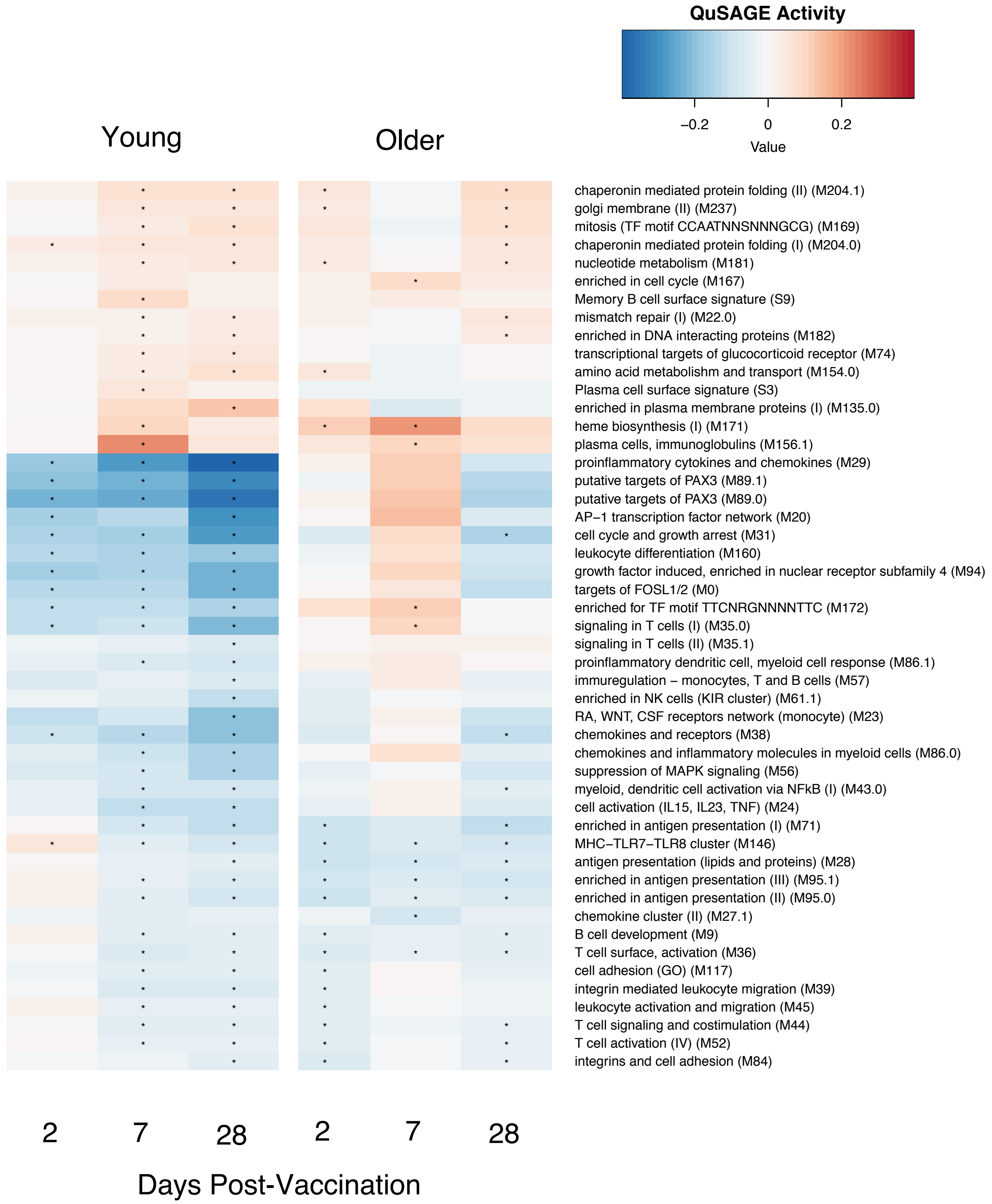


**Fig. S6. Comparison of Consistent Vaccine-Induced Changes in Gene Modules between Young and Older Adults.**

A heatmap of gene module meta-analysis activities for up- and down-regulated modules. Asterisks within the heatmap indicate modules significantly differentially expressed (FDR < 0.05) compared to day 0. Modules with FDR > 0.0025 or without annotation were removed for visualization. The full list of modules is available in *SI File 4*.

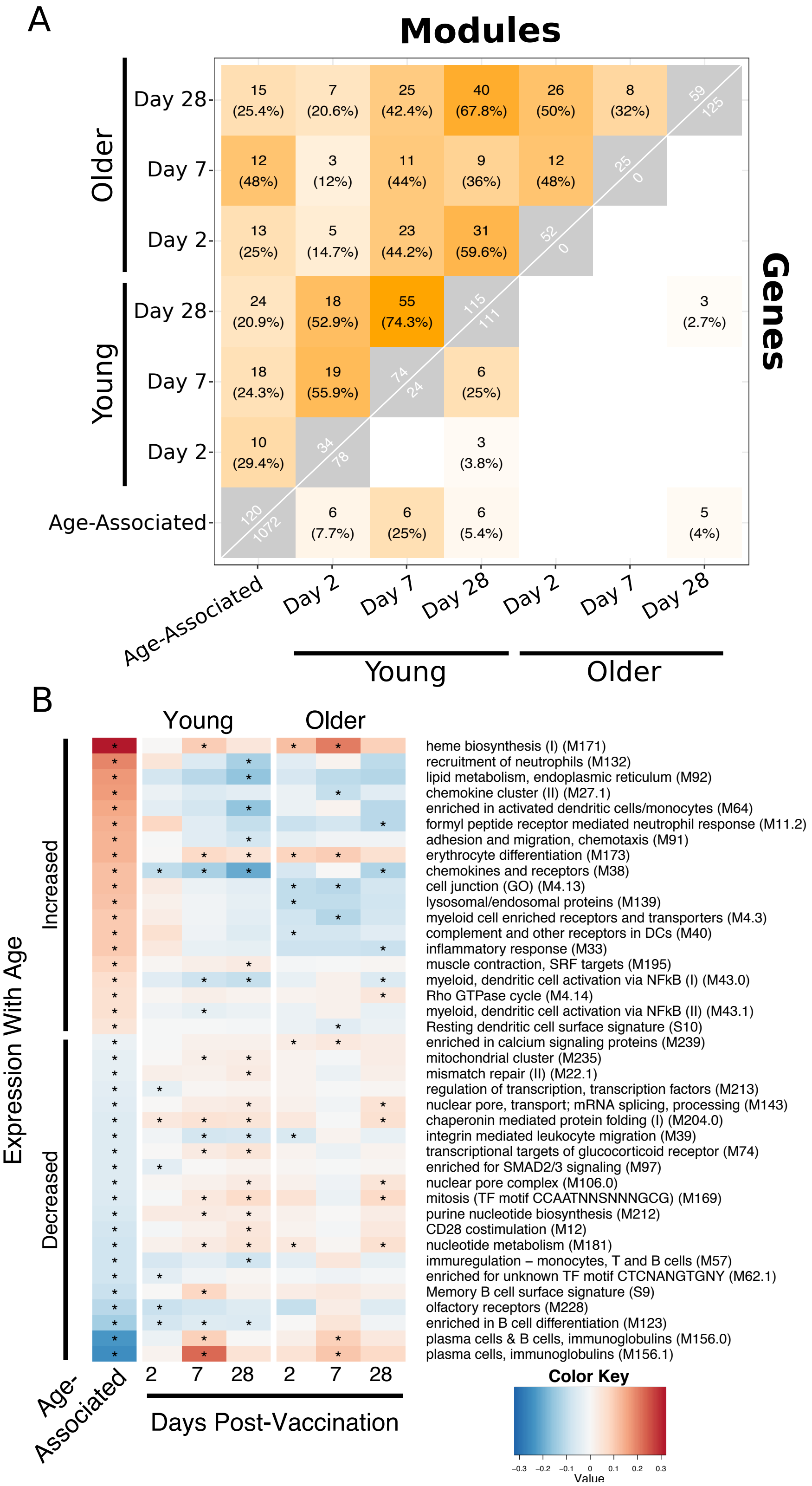
A
Modules
Genes
Older
Young
Day 28
Day 7
Day 2
Age-Associated
B
Expression With Age
Increased
Decreased
Young
Older
heme biosynthesis (I) (M171)
recruitment of neutrophils (M132)
lipid metabolism, endoplasmic reticulum (M92)
chemokine cluster (II) (M27.1)
enriched in activated dendritic cells/monocytes (M64)
formyl peptide receptor mediated neutrophil response (M11.2)
adhesion and migration, chemotaxis (M91)
erythrocyte differentiation (M173)
chemokines and receptors (M38)
cell junction (GO) (M4.13)
lysosomal/endosomal proteins (M139)
myeloid cell enriched receptors and transporters (M4.3)
complement and other receptors in DCs (M40)
inflammatory response (M33)
muscle contraction, SRF targets (M195)
myeloid, dendritic cell activation via NFkB (I) (M43.0)
Rho GTPase cycle (M4.14)
myeloid, dendritic cell activation via NFkB (II) (M43.1)
Resting dendritic cell surface signature (S10)
enriched in calcium signaling proteins (M239)
mitochondrial cluster (M235)
mismatch repair (II) (M22.1)
regulation of transcription, transcription factors (M213)
nuclear pore, transport; mRNA splicing, processing (M143)
chaperonin mediated protein folding (I) (M204.0)
integrin mediated leukocyte migration (M39)
transcriptional targets of glucocorticoid receptor (M74)
enriched for SMAD2/3 signaling (M97)
nuclear pore complex (M106.0)
mitosis (TF motif CCAATNNSNNNGCG) (M169)
purine nucleotide biosynthesis (M212)
CD28 costimulation (M12)
nucleotide metabolism (M181)
immuregulation – monocytes, T and B cells (M57)
enriched for unknown TF motif CTCNANGTGNY (M62.1)
Memory B cell surface signature (S9)
olfactory receptors (M228)
enriched in B cell differentiation (M123)
plasma cells & B cells, immunoglobulins (M156.0)
plasma cells, immunoglobulins (M156.1)
Age-Associated
2 7 28 2 7 28
Days Post-Vaccination
Color Key
Value

**Fig. S7. Age-Associated Signatures are Induced 7 Days Post-Vaccination**

(A) A heatmap of the overlap in age-associated and vaccine-induced genes or modules. Numbers in grey boxes along the diagonal show the number of genes (lower triangle) or modules (upper triangle) significantly altered in the specified condition. The off-diagonal boxes show the number of overlapping genes (lower triangle) or modules (upper triangle) between two conditions and the percent overlap (using the smaller set as the denominator). (B) A heatmap of gene modules that are significantly altered with aging and induced 2, 7, or 28 days post-vaccination. Asterisks within the heatmap indicate modules significantly differentially expressed compared to day 0 or between age groups, as indicated. Modules without annotation were removed for visualization (full list in *SI File 7*).

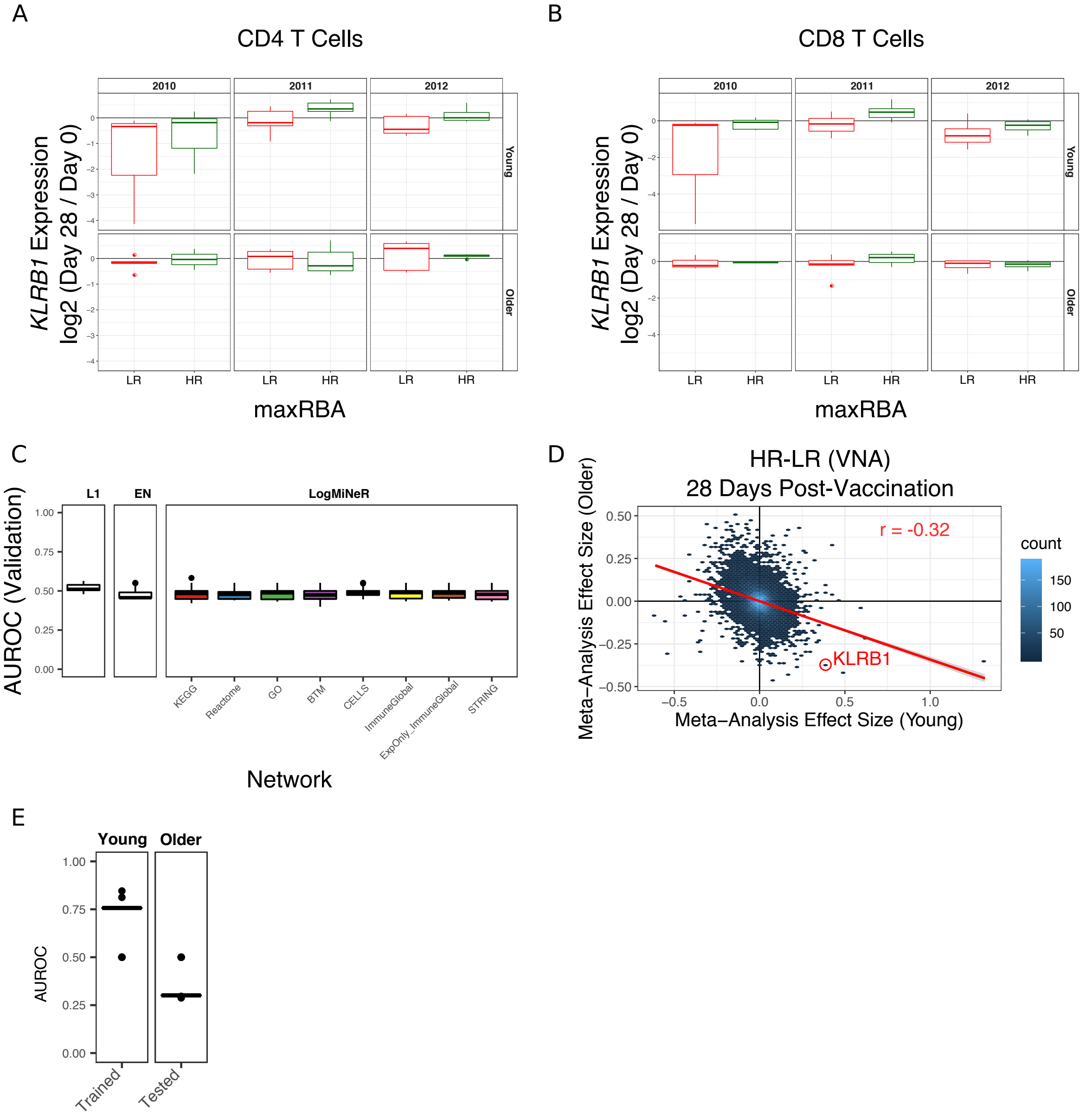


**Fig. S8. KLRB1 Expression in T Cells and Validation of Baseline Older Adult Models. Related to Figs. 4 and 5**

Boxplots of *KLRB1* expression changes 28 days post-vaccination in CD4 (A) and CD8 (B) T cells colored by response status (low responders (LR) or high responders (HR)) related to Fig. 4E. (C) Boxplots of the area under the receiver operating characteristic curve (AUROC) in the validation data (24) for models built from baseline transcriptional profiles in older adults. 50 iterations of cross-validation were performed. Related to Fig. 5B. (D) A scatter plot of the gene effect sizes comparing HR to LR (defined by virus neutralization assay, VNA) 28 days post-vaccination in young vs older adults. *KLRB1* is indicated as a gene that has a positive effect size in one age group and negative effect size in the other. Related to Fig. 4F. (E) A boxplot of the discovery and validation AUROC for models built on day 28 transcriptional profiles in young adults and tested on older adults. Related to Fig. 4E.

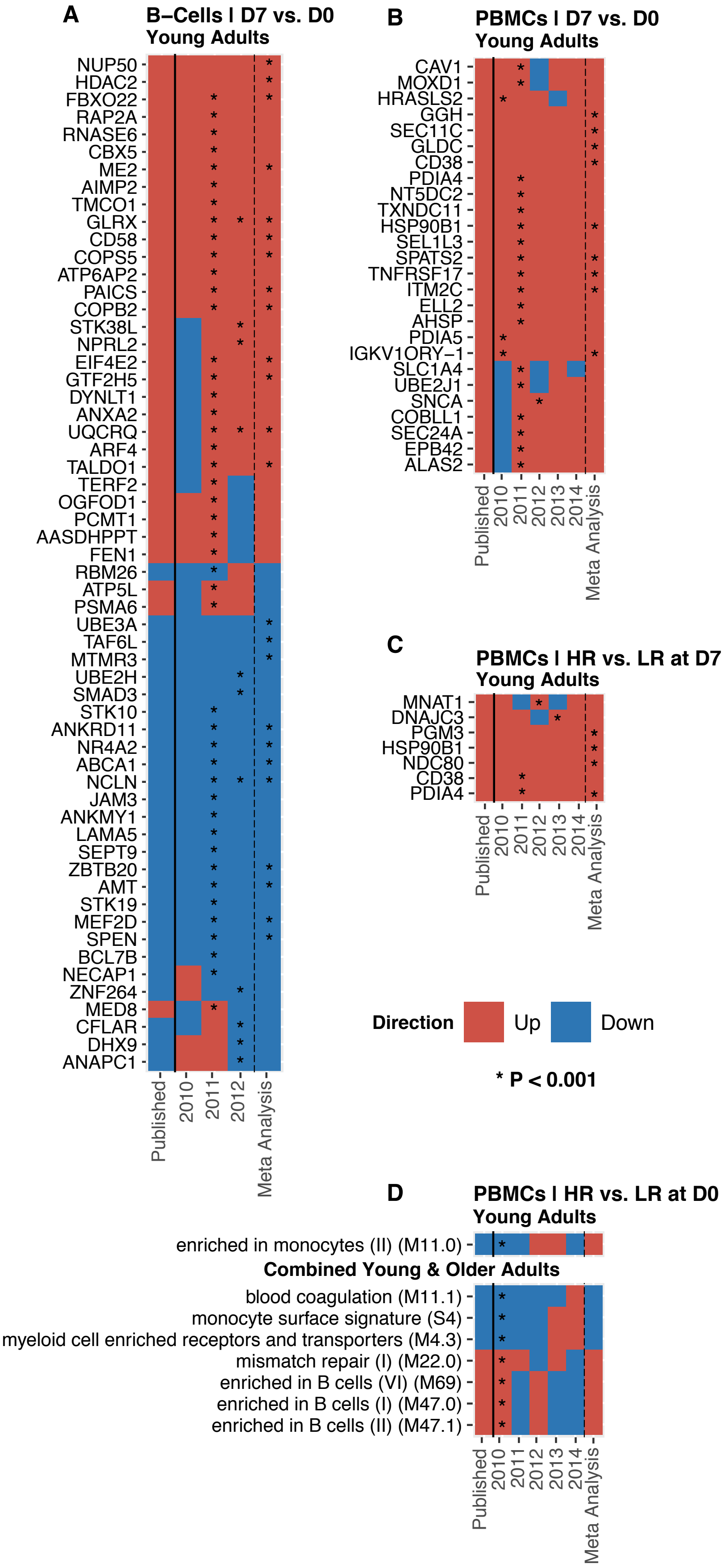
A
B–Cells I D7 vs. D0
Young Adults
NUPS50
HDAC2
FBXO22
RAP2A
RNASE6
CBX5
ME2
AIMP2
TMCO1
GLRX
CD58
COPS5
ATP6AP2
PAICS
COPB2
STK38L
NPRL2
EIF4E2
GTF2H5
DYNLT1
ANXA2
UQCRQ
ARF4
TALDO1
TERF2
OGFOD1
PCMT1
AASDHPPT
FEN1
RBM26
ATP5L
PSMA6
UBE3A
TAF6L
MTMR3
UBE2H
SMAD3
STK10
ANKRD11
NR4A2
ABCA1
NCLN
JAM3
ANKMY1
LAMA5
SEPT9
ZBTB20
AMT
STK19
MEF2D
SPEN
BCL7B
NECAP1
ZNF264
MED8
CFLAR
DHX9
ANAPC1
Published
2010
2011
2012
Meta Analysis
B
PBMCs I D7 vs. D0
Young Adults
CAV1
MOXD1
HRASLS2
GGH
SEC11C
GLDC
CD38
PDIA4
NT5DC2
TXNDC11
HSP90B1
SEL1L3
SPATS2
TNFRSF17
ITM2C
ELL2
AHSP
PDIA5
IGKV1ORY−1
SLC1A4
UBE2J1
SNCA
COBLL1
SEC24A
EPB42
ALAS2
Published
2010
2011
2012
2013
2014
Meta Analysis
C
PBMCs I HR vs. LR at D7
Young Adults
MNAT1
DNAJC3
PGM3
HSP90B1
NDC80
CD38
PDIA4
Published
2010
2011
2012
2013
2014
Meta Analysis
Direction
Up
Down
* P < 0.001
D
PBMCs I HR vs. LR at D0
Young Adults
enriched in monocytes (II) (M11.0)
Combined Young & Older Adults
blood coagulation (M11.1)
monocyte surface signature (S4)
myeloid cell enriched receptors and transporters (M4.3)
mismatch repair (I) (M22.0)
enriched in B cells (VI) (M69)
enriched in B cells (I) (M47.0)
enriched in B cells (II) (M47.1)
Published
2010
2011
2012
2013
2014
Meta Analysis

**Fig. S9. Many Published Signatures Show Similar Behaviors Over Multiple Seasons**
The expression levels of published gene signatures were analyzed to test whether they were consistently up (red) or down (blue) regulated. Transcriptional signatures from B cells comparing D7 vs. D0 (A), PBMCs comparing D7 vs. D0 (B), and PBMCs comparing the fold-changes (D7 vs. D0) in HR vs. LR (C) are shown as heatmaps. (D) Gene module signatures from PBMCs comparing the D0 expression in HR vs. LR. Analyses were context-matched based on cohort age. Young and older subjects are combined for some analyses as indicated. Asterisks indicate $p < 0.001$. The full results for all tested signatures are available in *SI File 10*.

**References**


1. Thakar J, et al. (2015) Aging-dependent alterations in gene expression and a mitochondrial signature of responsiveness to human influenza vaccination. *Aging (Albany NY)* 7(1):38–52.
2. van Duin D, et al. (2007) Prevaccine Determination of the Expression of Costimulatory B7 Molecules in Activated Monocytes Predicts Influenza Vaccine Responses in Young and Older Adults. *J Infect Dis* 195(11):1590–1597.
3. Steel J, et al. (2008) Live Attenuated Influenza Viruses Containing NS1 Truncations as Vaccine Candidates against H5N1 Highly Pathogenic Avian Influenza. *J Virol* 83(4):1742–1753.
4. Bentebibel S, et al. (2013) Induction of ICOS+CXCR3+CXCR5+ TH cells correlates with antibody responses to influenza vaccination. *Sci Transl Med* 5(176):176ra32.
5. Avey S, et al. (2017) Multiple network-constrained regressions expand insights into influenza vaccination responses. *Bioinformatics* 33(14):i208–i216.
6. Zapata HJ, et al. (2018) Impact of Aging and HIV Infection on the Function of the C-Type Lectin Receptor MINCLE in Monocytes. *Journals Gerontol Ser A* 74(6):794–801.
7. Gentleman R, et al. (2004) Bioconductor: open software development for computational biology and bioinformatics. *Genome Biol* 5(10):R80.
8. Yaari G, Bolen CR, Thakar J, Kleinstein SH (2013) Quantitative set analysis for gene expression: A method to quantify gene set differential expression including gene-gene correlations. *Nucleic Acids Res* 41(18):e170.
9. Ritchie ME, et al. (2015) limma powers differential expression analyses for RNA-sequencing and microarray studies. *Nucleic Acids Res* 43(7). doi:10.1093/nar/gkv007.
10. Kuleshov M V, et al. (2016) Enrichr: a comprehensive gene set enrichment analysis web server 2016 update. *Nucleic Acids Res* 44(W1):W90–W97.
11. Benjamini Y, Hochberg Y (1995) Controlling the False Discovery Rate: A Practical and Powerful Approach to Multiple Testing. *J R Stat Soc* 57(1):289–300.
12. Meng H, Yaari G, Bolen CR, Avey S, Kleinstein SH (2019) Gene set meta-analysis with Quantitative Set Analysis for Gene Expression (QuSAGE). *PLOS Comput Biol* 15(4):e1006899.
13. Viechtbauer W (2010) Conducting Meta-Analyses in R with the metafor Package. *J Stat Softw* 36(3):1–48.
14. Viechtbauer W (2005) Bias and efficiency of meta-analytic variance estimators in the random-effects model. *J Educ Behav Stat* 30(3):261–293.
15. Croft D, et al. (2014) The Reactome pathway knowledgebase. *Nucleic Acids Res* 42(D1):472–477.
16. Milacic M, et al. (2012) Annotating cancer variants and anti-cancer therapeutics in Reactome. *Cancers (Basel)* 4(4):1180–1211.
17. Ashburner M, et al. (2000) Gene Ontology: tool for the unification of biology. *Nat Genet* 25(May):25–29.
18. Li S, et al. (2014) Molecular signatures of antibody responses derived from a systems biology study of five human vaccines. *Nat Immunol* 15(2):195–204.
19. Abbas a R, et al. (2005) Immune response in silico (IRIS): immune-specific genes identified from a compendium of microarray expression data. *Genes Immun* 6(4):319–331.
20. Zhang JD, Wiemann S (2009) KEGGgraph: A graph approach to KEGG PATHWAY in R and bioconductor. *Bioinformatics* 25(11):1470–1471.
21. Gorenshteyn D, et al. (2015) Interactive Big Data Resource to Elucidate Human Immune Pathways and Diseases. *Immunity* 43(3):605–614.
22. Jensen LJ, et al. (2009) STRING 8 - A global view on proteins and their functional interactions in 630 organisms. *Nucleic Acids Res* 37(SUPPL. 1):412–416.

23. Tsang JS, et al. (2014) Global analyses of human immune variation reveal baseline predictors of postvaccination responses. *Cell* 157(2):499–513.
24. Nakaya HI, et al. (2015) Systems Analysis of Immunity to Influenza Vaccination across Multiple Years and in Diverse Populations Reveals Shared Molecular Signatures. *Immunity* 43(6):1186–1198.
25. Furman D, et al. (2013) Apoptosis and other immune biomarkers predict influenza vaccine responsiveness. *Mol Syst Biol* 9(659):659.
26. Furman D, et al. (2017) Expression of specific inflammasome gene modules stratifies older individuals into two extreme clinical and immunological states. *Nat Med* (January). doi:10.1038/nm.4267.
27. HIPC-CHI Signatures Project Team T, HIPC-I Consortium T (2017) Multicohort analysis reveals baseline transcriptional predictors of influenza vaccination responses. *Sci Immunol* 2(14):eaal4656.
28. Nakaya HI, et al. (2011) Systems biology of vaccination for seasonal influenza in humans. *Nat Immunol* 12(8):786–795.
